# Discovery and remodeling of *Vibrio natriegens* as a microbial platform for efficient formic acid biorefinery

**DOI:** 10.1101/2022.12.15.520533

**Authors:** Jinzhong Tian, Wangshuying Deng, Ziwen Zhang, Jiaqi Xu, Guoping Zhao, Sheng Yang, Weihong Jiang, Yang Gu

## Abstract

Formic acid (FA), an organic one-carbon source that is easily produced from CO_2_, has emerged as a promising CO_2_-equivalent feedstock for one-carbon biorefinery. However, developing efficient formatotrophs for an economically competitive FA utilization system remains a grand challenge. Here, we discovered that the gram-negative bacterium *Vibrio natriegens* has exceptional FA tolerance and metabolic capacity natively. This strain was then remodeled by rewiring the serine cycle and the TCA cycle which resulted in a non-native closed loop (S-TCA) for enhancing FA assimilation. The rational design generated a powerful metabolic sink that enabled rapid emergence of the evolved strains with further significantly improved performance in using FA as the major or sole carbon source. Furthermore, the introduction of a foreign indigoidine-forming pathway into the best-performing *V. natriegens* strain (S-TCA-2.0) led to production of 29.0 g·L^-1^ indigoidine and consumption of 233.7 g·L^-1^ formate within 72 h, achieving an order of magnitude higher formate consumption rate (3.2 g·L^-1^·h^-1^) than the reported highest level in microorganisms. This work represents a significant step towards the development of industrially viable microorganisms for FA biorefinery.

---

Microbial assimilation and conversion of one-carbon (C1) compounds to produce value-added products have attracted significant attention because it represents a new and important biological manufacturing route using abundant and highly available carbon sources^1^. Formic acid (FA) is a promising C1 feedstock since it can be efficiently derived from CO_2_ or syngas (mainly CO and CO_2_) with high selectivity via chemical conversions and further utilized by microorganisms^2^. Additionally, the storage of FA as a liquid is more convenient compared to gases^3^. Thus, FA is a CO_2_-equivalent carbon source that is more easily handled than CO_2_.

Native formate-utilizing microorganisms mainly employ the Wood-Ljungdahl pathway (also known as the reduced acetyl-CoA pathway), serine cycle and reductive glycine pathway for FA assimilation^4^. These pathways all depend on a core FA-fixing module called the tetrahydrofolate (THF) cycle which is initiated by formate-tetrahydrofolate ligase (FTL)^4^. Additionally, pyruvate formate-lyase (PFL)-catalyzed pyruvate formation from formate and acetyl-CoA is another important formate-fixing reaction in organisms^4^. However, native formate-utilizing microorganisms generally grow slowly and cannot efficiently utilize FA^5–17^. Hence, efforts have been made to develop synthetic formatotrophs, such as *Escherichia coli*, through rational genetic modifications and adaptive laboratory evolution, wherein FA assimilation was achieved by reconstructing the THF cycle and reverse glycine cleavage^18–20^. Despite the improved performances of these engineered *E. coli* strains in FA utilization, the efficiency is still much lower than those of traditional carbon sources such as sugars, which is largely due to the native weak FA tolerance and metabolic capacity of *E. coli*. Therefore, the discovery and employment of more suitable microorganisms as a chassis to establish efficient FA bioconversion platforms are needed.

*Vibrio natriegens* is a rapidly growing bacterium with the shortest doubling time (< 10 min) of all known bacteria^21,22^. It is also regarded as a desirable microbial host for bio-manufacturing^23–25^. This bacterium is able to grow on diverse substrates including galactose, arabinose, and glycerol^26,27^, probably due to its versatile metabolism. Furthermore, genetically modified *V. natriegens* strains can produce multiple chemicals including amino acids, polymers and polyols^27–29^. However, its inherent physiological and metabolic characteristics and potential in utilizing C1 resources remain unreported. This study surprisingly discovered that *V. natriegens* possesses superior tolerance and metabolic capacity to FA. The bacterium was further designed and modified into an exceptional microbial platform for FA utilization and conversion.

## Results

### Native *Vibrio natriegens* has superior formate tolerance and metabolic capacity

*V. natriegens* has rapid growth, but its formate tolerance and metabolic capacity are unknown. Thus, we examined its growth and formate consumption in the presence of different formate concentrations. *V. natriegens* could grow in the LBv2 medium supplemented with 20, 40 and 60 g·L^-1^ sodium formate (HCOONa·2H_2_O) (equivalent to 192, 385 and 577 mM formate, respectively), and consumed 186, 286 and 148 mM formate respectively within 24 h, although the growth rates and biomass gradually decreased with increasing formate concentration (Fig. 1a). However, at the higher sodium formate concentration (80 g·L^-1^, equivalent to 769 mM formate), the strain could not grow, largely due to the tolerance limit (Fig. 1a). A comparison with the data reported in representative formate-utilizing microorganisms showed that *V. natriegens* has the best formate tolerance and metabolic ability (Supplementary Table 1), showcasing great potential in biological FA utilization.

**Fig. 1.**
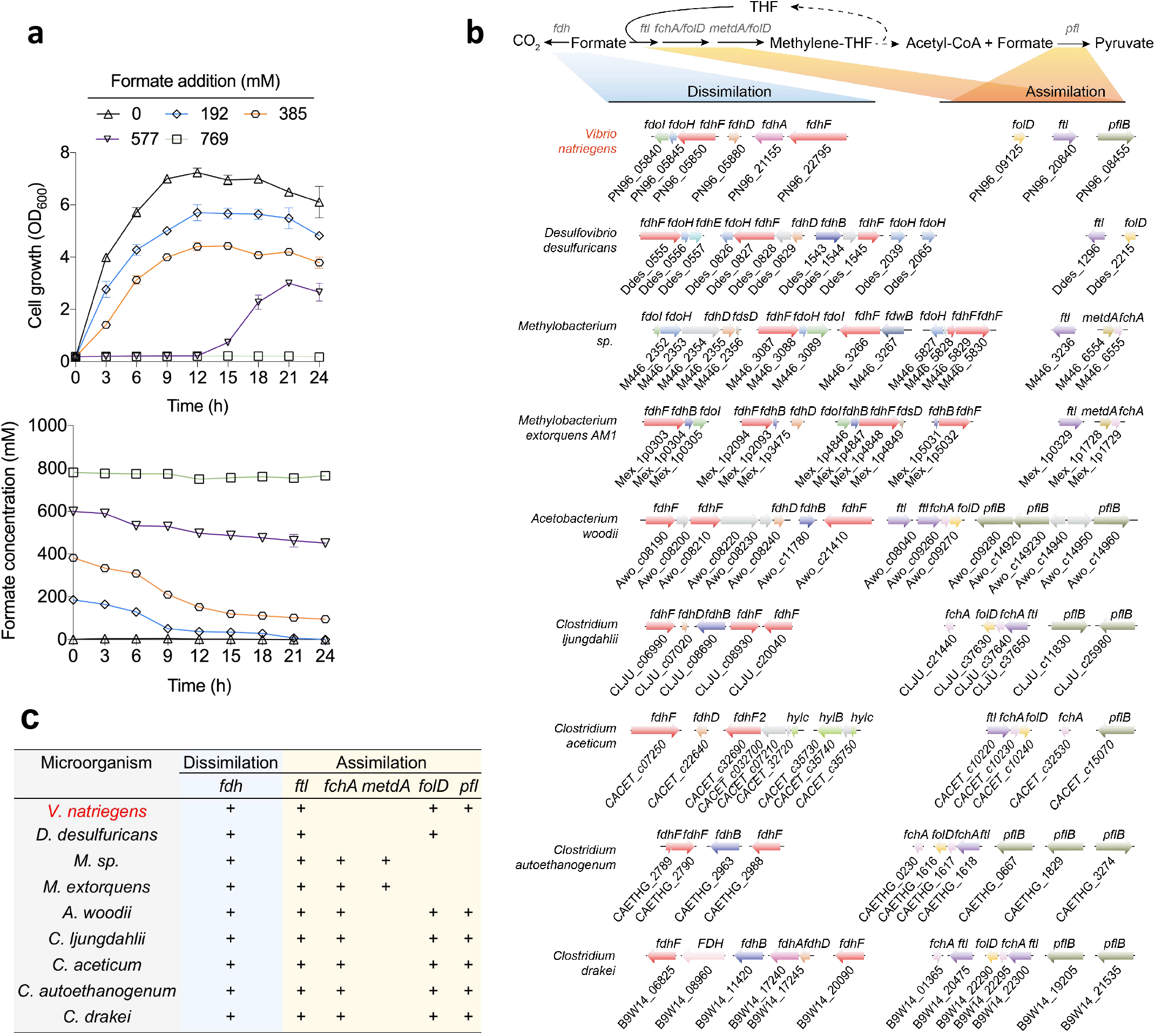
Comparative analysis of formate metabolic pathways in *V. natriegens*. **a,** Cell growth and formate consumption profiles with the supplementation of different amounts of sodium formate (HCOONa·2H_2_O). Data are presented as the mean ± SD (n = 3). Error bars show SDs. **b,** Putative genes and gene clusters involved in formate assimilation and dissimilation pathways in *V. natriegens* and some other representative formate-utilizing microorganisms. Homologous genes are marked with the same colors. *fdoI:* formate dehydrogenase subunit gamma; *fdhB:* formate dehydrogenase (NADP^+^) beta subunit; *fdoH:* formate dehydrogenase iron-sulfur subunit; *fdhF:* formate dehydrogenase major subunit alpha; *fdsD*: formate dehydrogenase subunit delta; *fdhA*: formate dehydrogenase (coenzyme F420) alpha subunit; *fdwB:* formate dehydrogenase beta subunit; *FdhD*: formate dehydrogenase accessory protein; *ftl*: formate--tetrahydrofolate ligase; *fchA*: methenyltetrahydrofolate cyclohydrolase; *mtdA*: Methylenetetrahydrofolate dehydrogenase (NADP^+^); *folD*: bifunctional methylenetetrahydrofolate dehydrogenase/methenyltetrahydrofolate cyclohydrolase; *pflB*: pyruvate formate-lyase. **c**, Occurrence of genes involved in formate metabolism (assimilation and dissimilation) in *V. natriegens* and some other representative formate-utilizing microorganisms. The presence of genes for the corresponding functions is shown with a “+”. *V. natriegens: Vibrio natriegens; D. desulfuricans: Desulfovibrio desulfuricans; M. sp.: Methylobacterium species; M. extorquens: Methylobacterium extorquens A. woodii: Acetobacterium woodii; C. ljungdahlii: Clostridium ljungdahlii; C. aceticum: Clostridium aceticum; C. autoethanogenum: Clostridium autoethanogenum; C. drakai: Clostridium drakai*.

### Genes and metabolic pathways for formate metabolism in *V. natriegens*

Natural FA-fixing reactions in organisms are quite scarce, in which two of the best known are FTL-catalyzed formylation of tetrahydrofolate (THF) and (PFL)-catalyzed pyruvate formation (Supplementary Fig. 1). Additionally, FA can be oxidized to form CO_2_ by formate dehydrogenase (FDH), a major FA dissimilation reaction in microorganisms. The *V. natriegens* genome contains three *fdh* genes (two *fdhFs* and one *fdhA*), two THF cycle genes (*ftl* and *folD*), and one *pfl* gene, enabling the operation of the abovementioned FA dissimilation and assimilation reactions in cells (Fig. 1b); however, the other two THF cycle genes, *fchA* and *metdA*, were missing (Fig. 1c). Furthermore, all of the genes responsible for the reactions starting from methylene-THF in the reductive glycine pathway and serine cycle were found in *V. natriegens* (Supplementary Fig. 2a), indicating that the assimilated formate can be converted into diverse metabolites. Energy conservation systems play important roles in microbial growth on organic C1 resources such as formate and methanol^20^. In the *V. natriegens* genome, diverse energy conservation systems were found, most of which had multiple coding genes and gene clusters (Supplementary Fig. 2b). This may be crucial for the required energy supply in the growth of *V. natriegens* on formate.

To explore functional genes and metabolic pathways for formate metabolism in *V. natriegens*, we measure the transcriptomic changes induced by formate using RNA-seq. There were 1,361 and 606 genes significantly up-regulated and down-regulated in *V. natriegens*, respectively, following formate addition to the culture (Supplementary Fig. 3a). Functional category analysis yielded 28 subsets containing many genes exhibiting significantly altered transcriptional levels (Supplementary Fig. 3b). An obvious positive correlation was detected between qRT-PCR and RNA-seq results for 15 selected genes (Supplementary Fig. 4), suggesting a good quality of the comparative transcriptomic data. We observed that, with formate stress, multiple crucial genes located in the THF cycle, the reductive glycine pathway and the serine cycle were significantly up-regulated. Two (PN96_05850 and PN96_22795) out of three FDH-encoding genes were up-regulated with a higher fold change (9.7-fold) occurring for the former one. Furthermore, a 8.9-fold transcriptional increase in the *pfl* gene encoding pyruvate formate-lyase occurred (Fig. 2a). These findings strongly suggest that the formate assimilation and dissimilation pathways mediated by these genes contribute to formate metabolism in *V. natriegens*. Noticeably, the transcriptional levels of nearly all the genes located in the TCA cycle were significantly up-regulated (Fig. 2a). The TCA cycle is the metabolic hub in cells connecting major nutrients, including sugars, lipids, and amino *acids*^30, 31^. The overall up-regulation of TCA genes indicates that *V. natriegens* enhanced its fundamental metabolism to meet the energy demand of formate utilization. In addition, multiple genes encoding ionic transporters or associated with unsaturated fatty acid synthesis and DNA repair were significantly up-regulated (Supplementary Fig. 5). Microbial acid tolerance can be affected by ionic transporter efficiency and the ratio of unsaturated fatty acids in cell membrane^32–34^. Hence, higher expression of these pathway genes may contribute to the formate tolerance of *V. natriegens*.

**Fig. 2.**
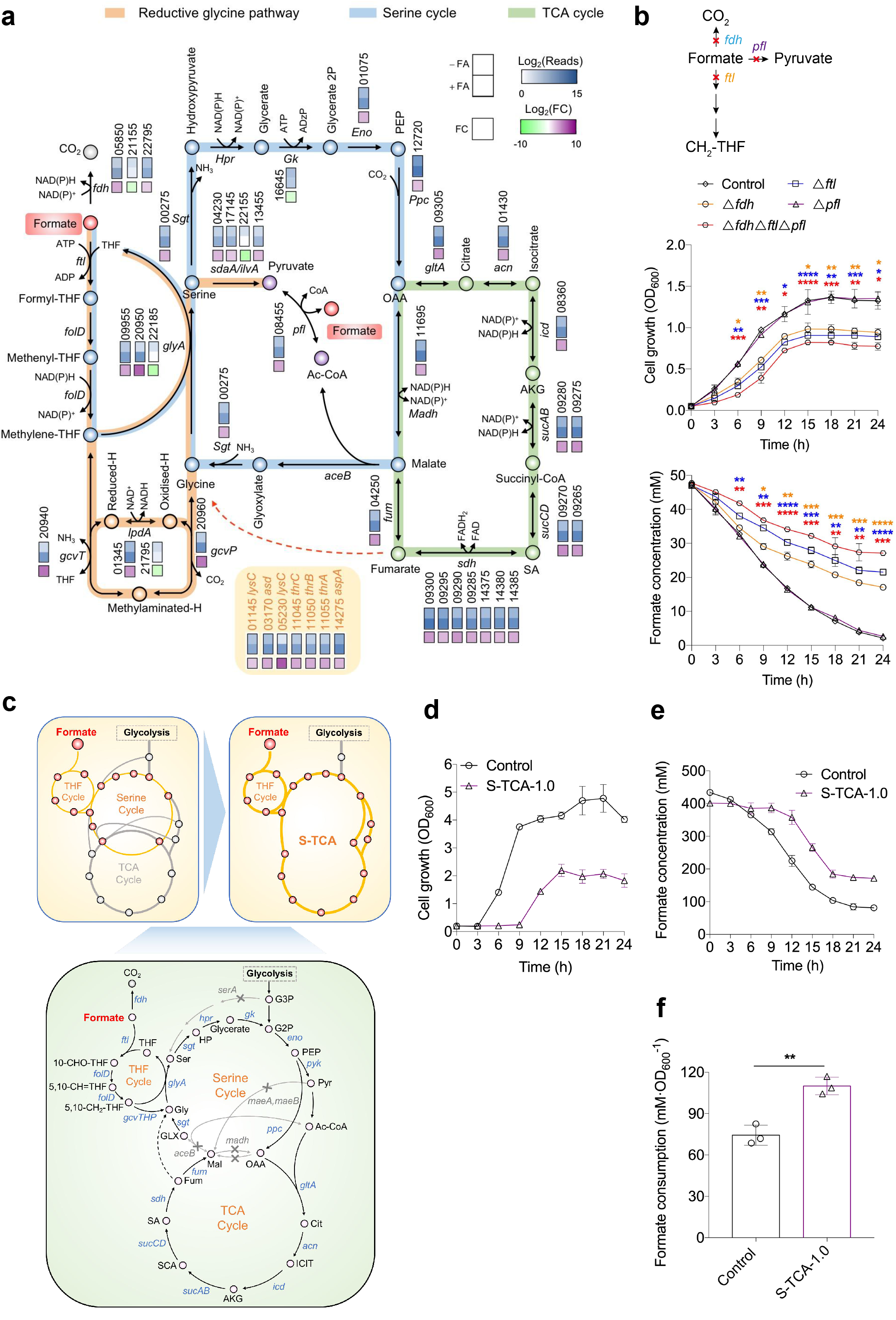
Validation of formate assimilation and dissimilation pathways and reconstitution of formate metabolism in *V. natriegens*. **a**, Transcriptional differences of genes involved in the tetrahydrofolate (THF) cycle, serine cycle, pyruvate formate-lyase (PFL) pathway, and formate dehydrogenase (FDH)-mediated formate dissimilation pathway with and without formic acid (FA) supplementation in *V. natriegens*. The bottom box indicates fold change as gene expression with FA over gene expression without FA. **b**, Influence of blocking the FTL-mediated THF cycle, pyruvate formate-lyase (PFL) pathway, and formate dehydrogenase (FDH)-mediated formate dissimilation pathway (upper) on cell growth (middle) and formate metabolism (bottom) of *V. natriegens*. The deleted *ftl* and *pfl* genes were PN96_20840 and PN96_08455, respectively. The *fdh-deleted* mutant was constructed by simultaneous deletion of the three annotated *fdh* genes (PN96_05850, PN96_21155, and PN96_22795). Sodium formate (HCOONa·2H_2_O) supplemented in the modified M9 medium was 5.0 g·L^-1^. Data are presented as the mean ± SD (*n* = 3). **c,** The combined manipulations fused the serine cycle and TCA cycle to creat a metabolic sink to promote formate metabolic flux. **d**, **e**, The growth (d) and total formate consumption (e) of the wild type and reconstructed (S-TCA-1.0) strains in the LBv2 medium supplemented with 40 g·L^-1^ sodium formate. **f**, Formate consumption per unit biomass of the wild type and S-TCA-1.0 strains in the LBv2 medium supplemented with 40 g·L^-1^ sodium formate. Data are presented as the mean ± SD (*n* = 3). Statistical analysis was performed by a two-tailed Student’s *t*-test. *, *P* < 0.05; **, *P* < 0.01; ***, *P* < 0.001 versus the control strain.

Based on the above analyses, *V. natriegens* is predicted to metabolize formate through FTL and PFL-catalyzed assimilation reactions and a FDH-catalyzed dissimilation reaction (Fig. 2b). To test them, the essential genes responsible for these reactions, i.e., *fdh, ftl* and *pfl*, were separately or simultaneously deleted in *V. natriegens* for phenotypic analysis (Fig. 2b, upper). The deletion of *fdh* or *ftl* significantly impaired cell growth and formate consumption, while the *pfl* deletion had no obvious influence (Fig. 2b, middle and bottom). Therefore, it seems that *fdh* and *ftl* mediated formate oxidation and assimilation, respectively, play important roles in formate metabolism in *V. natriegens*. Of note, although the simultaneous deletion of *fdh, ftl* and *pfl* further decreased formate consumption compared to the single *fdh* or *ftl* deletion, the strain with triple mutations maintained partial formate consumption ability (Fig. 2b, bottom), indicating that these reactions were not completely blocked or there are other unknown pathways responsible for formate metabolism in *V. natriegens*.

### Designing and creating a metabolic sink to promote formate metabolic flux

The above analyses prompted us to reprogram *V. natriegens* to further enhance its formate consumption. The following basic rationale was adopted: construction of a powerful metabolic sink that is linked to formate assimilation by integrating the TCA cycle and the serine cycle, thereby increasing the flux of the serine cycle and its upstream THF cycle by the strong pull from the TCA cycle (Fig. 2c, upper). This was achieved by manipulating the following pathways and genes (Fig. 2c, bottom): (i) disruption of three *madh* genes (PN96_06470, PN96_19465, PN96_11695), two *maeAB* genes (PN96_07295 and PN96_14755) and the *aceB* gene (PN96_10585) to fuse the serine cycle and the TCA cycle, forcing the metabolic flux of the serine cycle to go through the TCA cycle; (ii) disruption of the *serA* gene (PN96_00930) to enable more serine generation from formate assimilation rather than glycolysis.

The resulting mutated strain (S-TCA-1.0) was cultivated in media supplemented with 40 g·L^-1^ sodium formate (HCOONa·2H_2_O) (equivalent to 385 mM formate). Although S-TCA-1.0 exhibited impaired growth and less total formate consumption compared to the wild-type strain (Fig. 2d and e), its formate consumption per unit biomass was much higher than that of the latter (110 mM·OD_600_^-1^ versus 74 mM·OD_600_^-1^) (Fig. 2f). These results strongly indicate the effectiveness of our metabolic engineering strategy in enhancing the formate metabolic flux of *V. natriegens* cells, thereby laying the foundation for further strain improvement.

### Adaptive laboratory evolution of S-TCA-1.0 leads to high-performance strains with formate as the main or sole carbon source

Subsequently, the S-TCA-1.0 strain was subjected to adaptive laboratory evolution with gradually increasing sodium formate concentrations to improve its growth under formate stress, aiming to enhance total formate consumption. The strain was initially cultivated in the LBv2 medium supplemented with 30 g·L^-1^ sodium formate (288 mM formate) and subcultured once every 12 h (Fig. 3a). Several rounds of subculturing resulted in the biomass of grown cells reaching OD_600_ ~1.0. At this point, the sodium formate concentration was increased and a new evolutionary period began. This approach gradually improved S-TCA-1.0 tolerance to formate. Finally, one hundred and sixty-eight serial subculturing events generated an evolved strain (named S-TCA-2.0) capable of growing in the presence 150 g·L^-1^ sodium formate (1,442 mM formate) (Fig. 3b). Surprisingly, S-TCA-2.0 also exhibited exceedingly high formate utilization ability, capable of growing to OD_600_ ~1.6 with the consumption of 114.7 g·L^-1^ sodium formate (1,103 mM formate) within 24 h following initial supplementation of 125 g·L^-1^ sodium formate (1,202 mM formate) in the medium (Fig. 3c). An independent adaptive laboratory evolution strategy was performed in parallel to improve the formate utilization of the *V. natriegens* wild-type strain; however, the cellular adaptation of this strain to formate was much slower than that of S-TCA-2.0, and enhancement was difficult when the sodium formate concentration reached 100 g·L^-1^ (962 mM formate) (Supplementary Fig. 6). These results suggest that the reconstituted formate metabolic pathways in *V. natriegens* facilitated the strain evolution to adapt to high formate stress and improved formate metabolic capacity.

**Fig. 3.**
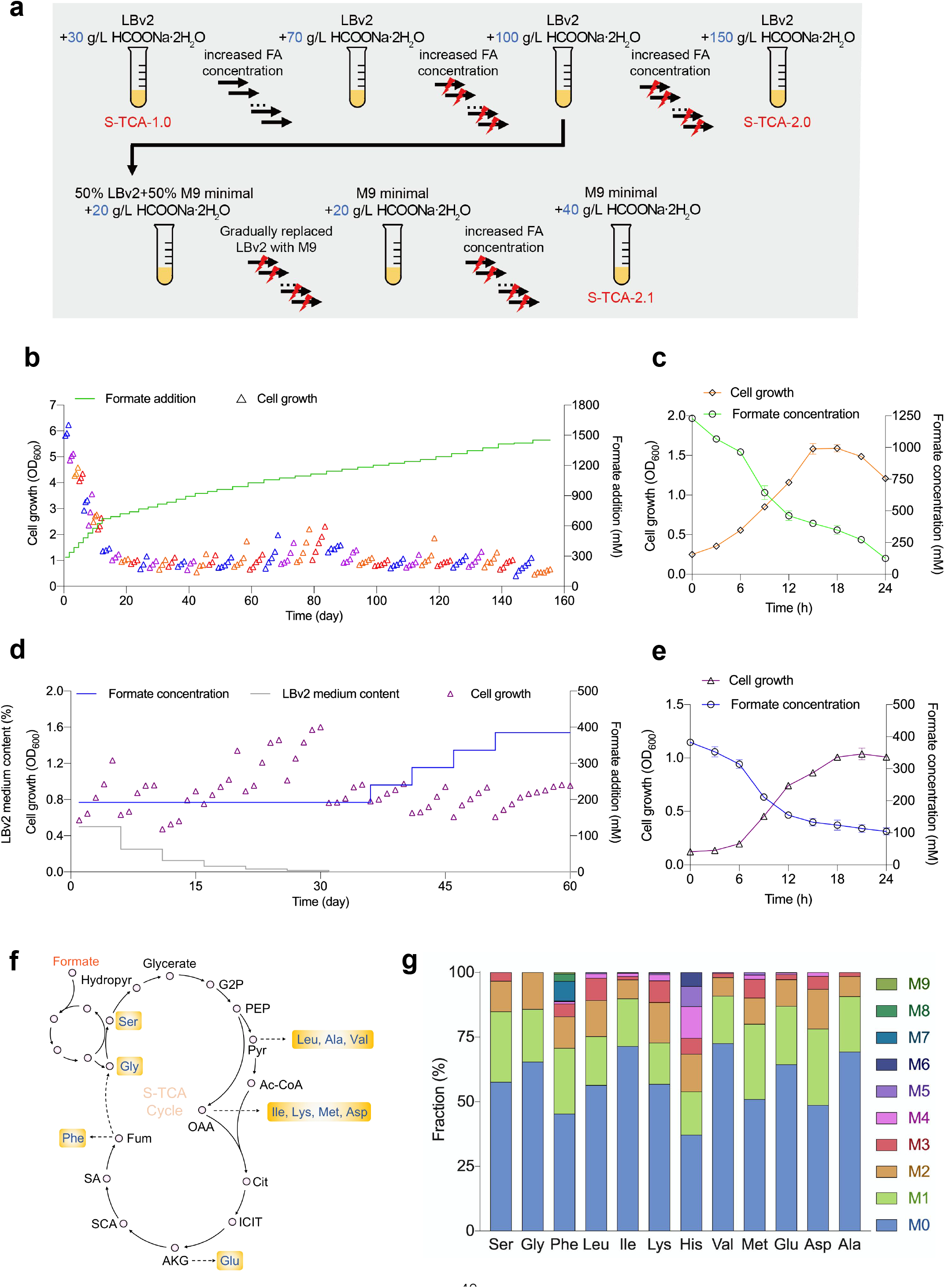
Adaptive laboratory evolution of the *V. natriegens* strains further enhances formate metabolism, or enables growth using formate as the sole carbon source. **a,** Flowchart for the adaptive laboratory evolution of the *V. natriegens* S-TCA-1.0. S-TCA-2.0 was the final evolved strain with significantly enhanced formate metabolism ability in LBv2 medium, while S-TCA-2.1 was the evolved strain capable of using formate as the sole carbon source. The lightning symbols represent UV mutagenesis, which was carried out every three passages, aiming to accelerate the strain mutation and evolutionary efficiency. **b**, Evolution trajectory of S-TCA-1.0 to obtain the S-TCA-2.0 strain with gradually enhanced formate utilization. LBv2 medium was used with an increasing sodium formate (HCOONa·2H_2_O) concentration from 30 to 150 g·L^-1^ (288 to 1,442 mM equivalent formate). Triangles indicate the bacterial biomass of each passage; the same colors represent passages with the same formate concentration. **c**, Validation of the growth and formate consumption of S-TCA-2.0 within 24 h using LBv2 medium initially supplemented with 125 g·L^-1^ sodium formate (1,202 mM equivalent formate). Data are presented as the mean ± SD (*n* = 3). **d**, Evolution trajectory of S-TCA-1.0 to obtain the S-TCA-2.1 strain capable of using formate as the sole carbon source. The medium consisted of gradually decreasing and increasing LBv2 and defined minimal M9 medium, respectively, until M9 was the only component. Triangles indicate the bacterial biomass of each passage; the same colors represent the passages with identical formate concentration. **e**, Validation of the growth and formate consumption of S-TCA-2.1 within 24 h using M9 medium initially supplemented with 40 g·L^-1^ sodium formate (385 mM equivalent formate). Data are presented as the mean ± SD (*n* = 3). **f**, Metabolic pathways synthesizing amino acids from formate in *V. natriegens*. Dashed lines represent multiple-step pathways. **g**, Ratios of ^13^C-labeled amino acids of the S-TCA-2.1 strain. Data are presented as the mean ± SD (*n* = 3).

Furthermore, we explored whether the reconstituted S-TCA-1.0 strain can evolve to use formate as the sole carbon source for growth. The initial subculturing process was the same as the adaptive laboratory evolution of S-TCA-1.0 leading to S-TCA-2.0. When the culture could grow on 100 g·L^-1^ sodium formate, it was transferred into a 1:1 mixture of the LBv2 and modified M9 media containing 192 mM formate for the following subculturing (Fig. 3a). The culture grew to OD_600_ ~ 1.2 within 24 h after five passages (Fig. 3d). The amount of the LBv2 medium was then sequentially decreased and replaced by the modified M9 medium until the culture could grow on formate as the sole carbon source (Fig. 3d). The final evolved strain was named S-TCA-2.1 and it could grow to OD_600_ ~1.0 within 24 h when cultivated in the modified M9 medium containing 40 g·L^-1^ sodium formate (385 mM formate), with consumption of 278 mM formate (Fig. 3e). These results demonstrate that we develope a *V. natriegens* strain capable of using formate as the sole carbon source.

For further confirmation, isotopic labeling experiments were performed for S-TCA-2.1. The strain was cultivated in the modified M9 medium containing both ^13^C-labeled formate and unlabeled formate (25% versus 75%) for continuous passage (three times), aiming to ensure that all the ^13^C-labeled metabolites reached a steady state. The following gas chromatography-mass spectrometry (GC-MS) showed that all the detected amino acids associated with the THF-cycle, serine cycle and reductive glycine pathway were labeled in cells, including serine, glycine, leucine, lysine, valine, alanine, glutamate, isoleucine, aspartate, phenylalanine, and methionine (Fig. 3f and g). Together, both formate fermentation experiments and isotopic labeling tests validate that S-TCA-2.1 can use formate as the sole carbon source to support its growth.

### Engineering the strain S-TCA-2.0 for efficient synthesis of indigoidine from formate

Since the S-TCA-2.0 strain exhibited excellent formate utilization, we set out to evaluate if it could be reprogrammed to efficiently convert formate into valuable compounds. Here, indigoidine was chosen as a target product because it is an important pigment widely used in food, medicine, and dyeing industries, and moreover, can be synthesized using α-ketoglutarate in the TCA cycle as the precursor. The indigoidine-producing pathway consisting of *Streptomyces lividans* Idgs and *Bacillus subtilis* Sfp which catalyze the conversion of glutamine to indigoidine was introduced into S-TCA-2.0 via an expression plasmid, yielding the S-TCA-2.0-IE strain (Fig. 4a).

**Fig. 4.**
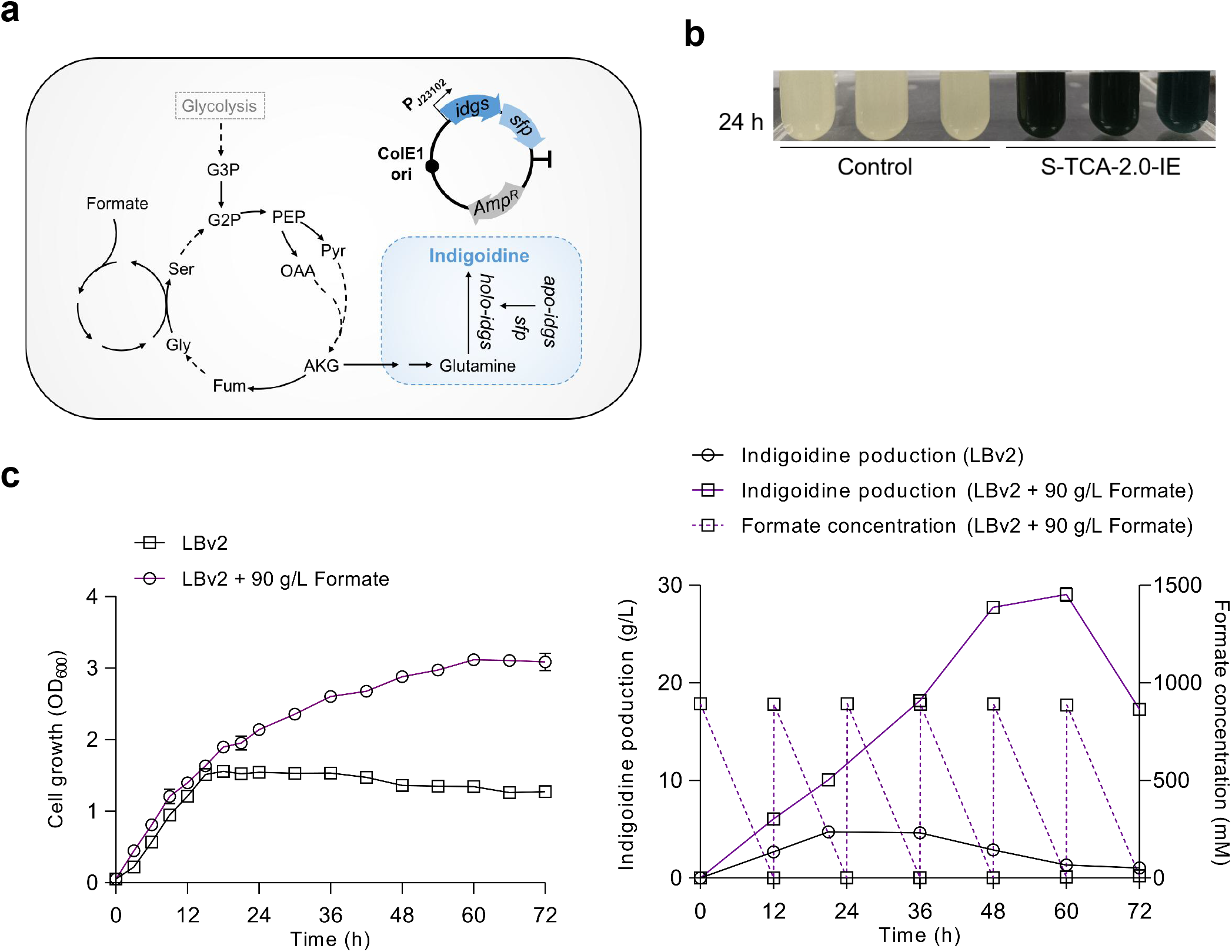
Engineering the evolved *V. natriegens* strain for efficient production of indigoidine from formate. **a**, The synthetic pathway and plasmid for indigoidine production. Idgs: indigoidine synthetase. Sfp: phosphoryltransferase. **b**, Indigoidine production by S-TCA-2.0-IE. The S-TCA-2.0 strain harboring a blank shuttle vector was used as the control. Strains were grown in LBv2 medium initially supplemented with 125 g·L^-1^ sodium formate (HCOONa·2H_2_O). **c**, Growth (left), indigoidine production (right), and formate consumption (right) of S-TCA-2.0-IE in a fed-batch mode. Initial sodium formate concentration in medium was 90 g·L^-1^. Extra sodium formate (90 g·L^-1^) was added into the medium every 12 h for five times. Data are presented as the mean ± SD (*n* = 3).

The S-TCA-2.0-IE strain was cultivated in the LBv2 medium supplemented with 125 g·L^-1^ sodium formate. As expected, the culture of S-TCA-2.0-IE appeared blue after 24 h of fermentation, whereas no blue occurred for the control (the S-TCA-2.0 strain harboring a blank plasmid), indicating that S-TCA-2.0-IE can produce indigoidine under high formate concentration (Fig. 4b). To explore the real potential of S-TCA-2.0-IE in converting formate into indigoidine, this strain was grown in the LBv2 medium containing 90 g·L^-1^ initial sodium formate, with additional sodium formate (90 g·L^-1^) added into the medium every 12 h (repeated five times), reaching to a total of 540 g·L^-1^ sodium formate supplementation (233.7 g·L^-1^ equivalent formate). S-TCA-2.0-IE continued to grow with formate supplementation, achieving the highest biomass (OD_600_ ~3.1) at 60 h (Fig. 4c, left); in contrast, the biomass of S-TCA-2.0-IE stopped to increase after 18 h in the medium without formate, reaching to the highest biomass of OD_600_ ~1.5 (Fig. 4c, left). Finally, S-TCA-2.0-IE consumed all the formate in the medium and produced 29.0 g·L^-1^ of indigoidine within 72 h, achieving a specific formate consumption rate of 3.2 g·L^-1^·h^-1^ (71.1 mM·h^-1^), which was an order of magnitude higher than the highest level reported in microorganisms (Supplementary Table 1). Meanwhile, indigoidine production of S-TCA-2.0-IE was only 4.7 g·L^-1^ without formate supplementation (Fig. 4c, right). These results demonstrate the excellent ability of S-TCA-2.0-IE to consume and convert formate, showcasing the potential of this engineered *V. natriegens* strain in FA biorefinery.

## Discussion

FA chemically converted from CO_2_ is an attractive feedstock for biological production of various value-added chemicals. Currently, all the known native formatotrophs are limited in practical application due to their low efficiency in using this C1 material. Genetically modified *V. natriegens* strains have been used to produce multiple products with high productivity by fermenting sugars and other cheap organic substrates^27^, indicating that this bacterium has exceptional substrate diversity and metabolic power. However, the potential of *V*. *natriegens* to utilize C1 materials has not been reported. In this study, we observed that multiple pathways and genes were activated in *V*. *natriegens* following high formate concentration stress; therefore, it seems that this organism has metabolic flexibility and versatility in the presence of this C1 substrate. Two out of three formate dehydrogenase genes were significantly up-regulated (Fig. 2a), indicating that these FDHs were activated to oxidize the available formate. Although the key genes (*ftl* and *folD*) responsible for the THF cycle exhibited similar expression levels with and without formate (Fig. 2a), the majority of genes associated with the downstream reductive glycine pathway, the serine cycle and the TCA cycle were significantly up-regulated following formate addition (Fig. 2a), which would pull the flux of the THF cycle and promote formate conversion into various metabolites.

Formate assimilation was previously used to relieve the serine-auxotrophy of *E. coli*^2, 35^, suggesting that the formate-assimilating flux could be directed toward serine synthesis. In our design, serine production from glycolysis was blocked by deleting the *serA* gene, which would enhance the THF cycle flux to form serine. Additionally, the artificial S-TCA cycle created by integrating the serine cycle and the TCA cycle enabled the high-flux TCA to pull the upstream carbon supply and enhance formate assimilation and conversion. This metabolic design is crucial for the following rapid evolution of *V*. *natriegens* into the strains with exceptional formate metabolic capacity since the wild-type strain did not achieve the goal using the same adaptive laboratory evolution strategy. Although the growth of *V*. *natriegens* was impaired after multiple gene deletions for the construction of S-TCA (Fig. 2d), adaptive laboratory evolution rapidly recovered cellular fitness. This finding indicates that *V*. *natriegens* has excellent plasticity in metabolic remodeling. Therefore, the rapid generation of the evolved strain (S-TCA-2.0) with strikingly enhanced formate metabolism should be attributed to the combination of metabolic engineering and adaptive laboratory evolution. This type of metabolic design may also work in other typical microbial chassis to create diverse cell factories for formate utilization.

It should be noted that formate assimilation to acetyl-CoA via the THF cycle in microorganisms is an energy consuming process. The energy for formate assimilation using formate as the sole carbon source mainly depends on FDH-catalysed formate oxidation which yields CO_2_ and leads to the loss of carbon. Moreover, all the reported native and artificial bacteria showed poor growth using formate as the sole organic carbon source^5–7, 9, 10, 19, 20^. Therefore, the LBv2 medium (containing yeast extract and peptone) was used here to evaluate the real potential of *V*. *natriegens* in formate consumption. Although yeast extract and peptone in the LBv2 medium will contribute to cell growth and indigoidine synthesis, formate consumption led to significantly higher biomass and indigoidine production of S-TCA-2.0-IE compared to the case without formate (Fig. 4c), suggesting efficient formate assimilation and its conversion to the target product in the strain. Therefore, the fermentation mode adopting formate as the primary carbon source supplemented with some other conventional substances as secondary carbon sources is likely a more feasible route for formate biorefinery currently compared to using formate as the sole carbon source. In summary, our work uncovered the excellent metabolic power of *V*. *natriegens* towards formate and further developed this bacterium for efficient formate conversion into target products. The discoveries and strategies presented here will effectively advance formate biorefinery.

## Methods

### Bacterial strains and growth conditions

*Vibrio natriegens* ATCC 14048 (stored at −80°C as a 20% glycerol stock) was used as the parental strain for formate metabolism as well as further genetic modifications. The *V. natriegens* strains were grown in the LBv2 nutrient-rich medium or a slightly modified M9 minimal medium (with addition of 2% (w/v) NaCl) at 30 °C^22^. Chloramphenicol (12.5 μg·mL^-1^), kanamycin (12.5 μg·mL^-1^), carbenicillin (100 μg·mL^-1^), and different amounts of sodium formate were added into the media when needed. *Escherichia coli* DH5α (stored at −20 °C as a 20% glycerol stock) was used as the host for plasmid cloning. The *E. coli* cells were grown in LB medium supplemented with chloramphenicol (12.5 μg·mL^-1^), ampicillin (100 μg·mL^-1^) or kanamycin (50 μg·mL^-1^). All the strains used in this study are listed in Supplementary Table 3.

### Reagents and chemicals

Primers were synthesized by BioSune (BioSune, Shanghai, China). PCR reactions were carried out by using KOD plus Neo and KOD FX DNA polymerase (Toyobo, Osaka, Japan). The restriction enzymes and ligases used in plasmid construction were purchased from Thermo Fisher Scientific (Thermo Fisher Scientific, Vilnius, Lithuania) and Takara (Takara, Dalian, China), respectively. The assembly of multiple DNA fragments for plasmid construction was performed by using the ClonExpress MultiS One Step Cloning Kit (Vazyme Biotech Co., Ltd., Nanjing, China). Plasmid isolation and DNA purification were performed with kits (Axygen Biotechnology Company Limited, Hangzhou, China).

^13^C-labelled sodium formate was ordered from Cambridge Isotope Laboratories (Cambridge Isotope Laboratories, lnc., MA, USA). The other commercial chemicals were purchased from Sigma-Aldrich (Sigma-Aldrich Co., St Louis, USA) and Aladdin (Shanghai Aladdin Biochemical Technology Co., Ltd., Shanghai, China). Sodium formate used in this study was HCOONa·2H_2_O.

### Construction of plasmids

All the plasmids and primers used in this study are listed in Supplementary Table 3 and Supplementary Table 4, respectively.

The vector co-expressing *idgs* (SLIVYQS_19313, encoding indigoidine synthetase) and *sfp* (GFX43_019515, encoding phosphoryltransferase), from *S. lavendulae* CGMCC 4.1386 and *Bacillus subtilis*, respectively, for indigoidine formation was constructed as follows. A large DNA fragment (*idgs-RBS-sfp*) that contained the codon-optimized *idgs* gene, an RBS sequence (ggatccactctgtcagatctcactctgccaaggaggacgcac), and the *sfp* gene was synthesized (GenScript, Nanjing, China). The other DNA fragment harboring a promoter (ttgacagctagctcagtcctaggtactgtgctagc) and RBS sequence (agagtcacacaggaaagtacta) were obtained by PCR amplification using the primers J23102-B00320m-F/J23102-B00320m-R, which have a 21-nt overlap between the 3’ region of the former primer and the 5’ region of the latter primer. Next, this DNA fragment was linked to the *idgs-RBS-sfp* fragment *via* overlapping PCR using the primers J23102-B00320m-idgs-sfp-F/idgs-sfp-R, yielding a larger DNA fragment. The yielded DNA fragment was further linked with a linear plasmid pColE1-Amp vector^36^ using a ClonExpress MultiS One Step cloning kit (Vazyme, Nanjing, China), generating the target plasmid for indigoidine formation.

The plasmid pMMB67EH-tfox-sacB, modified from the plasmid pMMB67EH-tfox^28^, was used for supporting the transformation of exogenous DNA into *V. natriegens*. In brief, the plasmid backbone was obtained by PCR amplification using the plasmid pMMB67EH-tfox as the template and the primers tfox-F/tfox-R. The *sacB* gene was obtained by PCR amplification using the plasmid pEcCas^37^ as the template and primers sacB-F/sacB-R. These two DNA fragments were assembled by the ClonExpress MultiS One Step cloning kit (Vazyme, China), yielding the plasmid pMMB67EH-tfox-sacB.

### Generation of *V. natriegens* mutants

Gene knockout was achieved in *V. natriegens* using the MuGENT (multiplex genome editing by natural transformation) method^28^. In brief, the *V. natriegens* cells harboring the abovementioned pMMB67EH-tfox-sacB plasmid were induced to competence by growing overnight in the LBv2 medium (containing 100 μg·mL^-1^ carbenicillin and 100 μM IPTG) in a shaking incubator with a shaker rate of 280 rpm at 30 °C. 3.5 μL of the grown culture were then diluted into 350 μL of the IO medium (28 g·L^-1^ of instant ocean sea salts)^28^ plus 100 μM IPTG, which was further mixed with 50 ng of a selected product (a DNA fragment that has an antibiotic resistance marker and two homologous arms flanking the *dns* gene) and 200 ng of an unselected product (a DNA fragment that has two homologous arms flanking the gene of interest but lacks any selectable marker). The mixture was incubated statically at 30 °C for 5 h. Next, 1 mL of the LBv2 medium was added, and the mixture was further outgrown at 30 °C with shaking (280 rpm) for 2 h. The cultures were then plated onto LBv2 agar plates (containing 100 μg·mL^-1^ carbenicillin and 12.5 μg·mL^-1^ chloramphenicol or 100 μg·mL^-1^ kanamycin). When colonies were visible on the plates after incubation at 30 °C for 10 h, some were picked and subjected to colony PCR using the primers KO-gene-seq-F/KO-gene-seq-R, aiming to detect the desired gene knockout event.

Finally, to eliminate the pMMB67EH-tfox-sacB plasmid in cells, the colonies with desired gene knockout were harvested from the agar surface, resuspended in 5 mL of the LBv2 medium (containing 80 g·L^-1^ sucrose), and incubated at 30 °C with shaking (280 rpm) for 8-10 h. The grown cells were plated onto the LBv2 agar plates (containing 80 g·L^-1^ sucrose) again. When colonies were visible on the plates after incubation at 30 °C for 12 h, some were picked and resuspended in the LBv2 medium with and without carbenicillin (100 μg·mL^-1^) to verify plasmid curing.

### Adaptive laboratory evolution

The engineered *V. natriegens* strain S-TCA-1.0 (0.1 mL) was first inoculated into 5 mL of the LBv2 medium containing 30 g·L^-1^ sodium formate and then incubated in a shaking incubator at 30 °C and 280 rpm. After overnight cultivation, 0.1 mL of grown cells were inoculated into 5 mL of the fresh LBv2 medium containing 30 g·L^-1^ sodium formate. Once cells grew to over OD_600_ 1.0 after overnight cultivation by serial subcultures, the sodium formate concentration was increased in the next round of subculturing. Samples were taken every passage and then stored at −80 °C. When formate concentration in the medium was raised to 70 g·L^-1^, the cultivation time of each passage strains were extended to 24 h, and simultaneously, the UV mutagenesis was carried out every three passages by placing the culture at a distance of 10 cm away from a 20-W UV light lamp for 15 min, aiming to accelerate the strain mutation and evolutionary efficiency.

When formate concentration in the medium reached to 100 g·L^-1^, the evolved cells were divided into two parts: one was further evolved in the LBv2 medium with gradually increased formate concentration; the other was evolved in 1: 1 of the LBv2: M9 (chemically defined minimal medium) media containing 20 g·L^-1^ of sodium formate, aiming to obtain an evolved *V. natriegens* strain capable of using formate as the sole carbon source. For the latter approach, when the cells could grow to OD_600_ ~1 within 24 h of cultivation, M9 ratio in the mixed medium was gradually increased to 75%, 87.5%, 93.75%, 96.87% and 98.43% for the following passages. Finally, LBv2 was completely eliminated from the medium, and an evolved *V. natriegens* strain (named as S-TCA-2.1) was obtained.

### Analytical methods

The concentration of formate was determined using an Agilent 1100 HPLC system (Agilent, Beijing, China) with a bio-Rad Aminex HPx-87H column (Bio-Red). In brief, samples were taken from fermentation broth at appropriate time intervals and then centrifuged at 7,000 × *g* for 20 min at 4 °C. Next, 200 μL of supernatant was analyzed by HPLC. The column temperature was maintained at 60 °C. The flow rate of the mobile phase (5mM H2SO4) is 0.6 mL·min^-1^.

The purification of indigoidine was performed with slight modifications as previously described^38^. In brief, the fermentation broth was freeze-dried. The generated powder was treated with three rounds of methanol, isopropanol, water, ethanol, and hexane, aiming to lyse cells and remove contaminating proteins and metabolites. The pellet was dried overnight and then resuspended in dimethylsulfoxide (DMSO) at a concentration of 1 mg·mL. Indigoidine purity was determined by the HPLC and mass spectrum (Supplementary Fig. 7). The purified indigoidine reagent was used to establish a standard curve correlating the indigoidine concentration to the absorbance at OD_600_ (Supplementary Fig. 8).

To quantify indigoidine produced by *V. natriegens*, 1 mL of the fermentation broth was collected and treated by the same procedure described above. The indigoidine concentration was determined by measuring the optical absorbance at 600 nm using a spectrophotometer (DU730, Beckman Coulter, Inc., USA).

### RNA-seq

*V. natriegens* strains were cultured in LBv2 medium with or without the supplementation of sodium formate (HCOONa·2H_2_O) (40 g·L^-1^). The grown cells (6 h of cultivation) were harvested by centrifugation (6,000 × *g*, 4 °C, 20 min) and then frozen in liquid nitrogen. The RNA-seq assays were finished by the Majorbio Co. Ltd. (Shanghai, China). In brief, total RNA was extracted using the TRIzol^®^ reagent (R0016, Beyotime, Shanghai) followed by the removal of genomic DNA using DNase I (TaKara). The 16S and 23S ribosomal RNA (rRNA) in total RNA were eliminated using the Ribo-Zero Magnetic kit (Epicenter Biotechnologies, WI, USA). Thereafter, mRNA was isolated using the polyA selection method with oligo (dT) beads, followed by fragmentation (100 to 400 bp) in fragmentation buffer. Next, cDNA was synthesized via reverse transcriptase using a SuperScript double-stranded cDNA synthesis kit (Invitrogen, CA) with random hexamer primers (Illumina). The second strand of cDNA was generated by incorporating deoxyuridine triphosphate (dUTP) into the place of deoxythymidine triphosphate (dTTP), aiming to create blunt-ended cDNAs. The yielding double-stranded cDNAs were subjected to end-repair, phosphorylation, and 3’ adenylation and adapter ligation in sequential. The second strand of cDNA with dUTP was degraded using UNG enzyme (Uracil-N-Glycosylase). The cDNA fragments of ~200 bp were excised from 2% Low Range Ultra Agarose gel. Then, cDNA libraries were obtained by PCR amplification for 15 cycles using Phusion DNA polymerase (NEB). After quantification with a micro fluorometer (TBS-380, TurnerBioSystems, USA), the libraries were sequenced on the Illumina HiSeq×Ten sequencer using paired-end sequencing. Bioinformatics analysis was performed using data generated by Illumina platform. All analyses were performed using a cloud platform of Shanghai Majorbio Bio-Pharm Technology Co., Ltd. The main methods are as follows: RSEM was used to quantify gene and isoform abundances. TPM method is used to calculated expression level. Differentially expressed genes were identified by using the DESeq2 packages (http://bioconductor.org/packages/release/bioc/html/DESeq2.html).

### qRT-PCR

qRT-PCR was performed to verify RNA-seq data. Briefly, cultures were taken at 6 h of cultivation in LBv2 medium with or without the supplementation of formate, frozen quickly in liquid nitrogen, and then ground into powder. RNA samples were extracted using the Ultrapure RNA Kit (SparkJade, Shanghai, China), following by the removal of genomic DNA with DNase I (SparkJade, Shanghai, China). Next, qRT-PCR was carried out in a MyiQ2 thermal cycler (Bio-Rad, USA) using a SYBR Green PCR premix kit (SparkJade, Shanghai, China). The relative transcript levels of each tested gene were normalized to the 16S rRNA gene (PN96_00205, an internal control) of *V. natriegens*. The primers used are listed in Supplementary Table 4.

### ^13^C isotopomer analysis

The *V. natriegens* cells were grown in the M9 medium at 30°C. 7.5 and 22.5 g·L^-1^ of ^13^C-labeled and unlabeled sodium formate, respectively, were added into the medium when the grown cells reached the plateau stage. After 24 h of cultivation, 0.01 mL of the grown cells were inoculated into the M9 medium (4 mL) for cultivation again with the supplementation of 7.5 and 22.5 g·L^-1^ of ^13^C-labeled and unlabeled sodium formate, respectively, at the plateau stage. Such a subculturing was repeated three times. The grown cells of the last passage were harvested by centrifugation (6,000 × *g*, 4°C, 20 min) and incubated with 200 μL 6M HCl at 105°C for 24 h. The samples were dried in a vacuum centrifuge and then derivatized with 100 μL of pyridine and 50 μL of *N*-methyl-*N*-[tert-butyldimethylsilyl] trifluoroacetamide at 85 °C for 1 h. After filtration (0.45 μm pore size, Millipore), 1 μL of samples were injected into a GC-MS system equipped with a DB-5HT column (30 × 0.25 mm, 0.1 μm), which operated in an electron shock (EI) mode at 70 eV. Amino acids were determined by matching masses and retention times to authenticated standards library.

## Data availability

RNA-seq data have been deposited in the NCBI SRA under bioproject ID PRJNA843433. All the data supporting the findings of this study are available in the text and the supplementary figures and tables or can be obtained from the corresponding authors upon reasonable requests.

## Acknowledgments

This research was supported by the National Key R&D Program of China (No. 2018YFA0901500), the National Natural Science Foundation of China (No. 31921006), the Shanghai Science and Technology Commission (No. 21DZ1209100), DNL Cooperation Fund, CAS (No. DNL202013), Tianjin Synthetic Biotechnology Innovation Capacity Improvement Project (TSBICIP-KJGG-016-01). We are grateful to Wenli Hu and Xiaoyan Xu for HPLC and LC-MS/MS analyses.

## Author contributions

J.T. conceived and performed strain modifications, adaptive laboratory evolution and isotopic labeling experiments; W.D. and Z.Z. constructed plasmids; J.T. and W.D. analyzed the phenotypic changes of the mutants; W.D. performed bioinformatics analyses; Y.G. and W.J. supervised and directed the study. Y.G., W.J., J.T. and W.D. wrote the manuscript. G. Z., S. Y. and J. X. discussed results and offered advices.

## References

1. Jiang, W. et al. Metabolic engineering strategies to enable microbial utilization of C1 feedstocks. Nat. Chem. Biol. 17, 845–855 (2021).

2. Yishai, O., Goldbach, L., Tenenboim, H., Lindner, S. N. & Bar-Even, A. Engineered assimilation of exogenous and endogenous formate in Escherichia coli. ACS Synth. Biol. 6, 1722–1731 (2017).

3. Bang, J. et al. Synthetic formatotrophs for one-carbon biorefinery. Adv. Sci. (Weinh) 8, 2100199 (2021).

4. Bar-Even, A. Formate assimilation: the metabolic architecture of natural and synthetic pathways. Biochemistry 55, 3851–3863 (2016).

5. Stokes, J. E. & Hoare, D. S. Reductive pentose cycle and formate assimilation in Rhodopseudomonas palustris. J. Bacteriol. 100, 890–894 (1969).

6. Goldberg, I. & Mateles, R. I. Growth of Pseudomonas C on C1 compounds: enzyme activites in extracts of Pseudomonas C cells grown on methanol, formaldehyde, and formate as sole carbon sources. J. Bacteriol. 122, 47–53 (1975).

7. Moon, Y. J. et al. Proteome analyses of hydrogen-producing hyperthermophilic archaeon Thermococcus onnurineus NA1 in different one-carbon substrate culture conditions. Mol. Cell. Proteomics 11, M111.015420 (2012).

8. Zhuang, W. Q. et al. Incomplete Wood-Ljungdahl pathway facilitates one-carbon metabolism in organohalide-respiring Dehalococcoides mccartyi. Proc. Natl Acad. Sci. USA 111, 6419–6424 (2014).

9. Grunwald, S. et al. Kinetic and stoichiometric characterization of organoautotrophic growth of Ralstonia eutropha on formic acid in fed-batch and continuous cultures. Microb. Biotechnol. 8, 155–163 (2015).

10. Urschel, M. R., Hamilton, T. L., Roden, E. E. & Boyd, E. S. Substrate preference, uptake kinetics and bioenergetics in a facultatively autotrophic, thermoacidophilic crenarchaeote. FEMS. Microbiol. Ecol. 92, fiw069 (2016).

11. Liu, Z. et al. Exploring eukaryotic formate metabolisms to enhance microbial growth and lipid accumulation. Biotechnol. Biofuels 10, 22 (2017).

12. Gonzalez de la Cruz, J., Machens, F., Messerschmidt, K. & Bar-Even, A. Core catalysis of the reductive glycine pathway demonstrated in Yeast. ACS Synth. Biol. 8, 911–917 (2019).

13. Arantes, A. L. et al. Enrichment of anaerobic syngas-converting communities and isolation of a novel carboxydotrophic Acetobacterium wieringae strain JM. Front. Microbiol. 11, 58 (2020).

14. Ergal, I. et al. Formate utilization by the crenarchaeon Desulfurococcus amylolyticus. Microorganisms 8, 454 (2020).

15. Sánchez-Andrea, I. et al. The reductive glycine pathway allows autotrophic growth of Desulfovibrio desulfuricans. Nat. Commun. 11, 5090 (2020).

16. Wood, G. E., Haydock, A. K. & Leigh, J. A. Function and regulation of the formate dehydrogenase genes of the methanogenic archaeon Methanococcus maripaludis. J. Bacteriol. 185, 2548–2554 (2003).

17. Lawson, C. E. et al. Autotrophic and mixotrophic metabolism of an anammox bacterium revealed by in vivo ^13^C and ^2^H metabolic network mapping. Isme j. 15, 673–687 (2021).

18. Bang, J. & Lee, S. Y. Assimilation of formic acid and CO2 by engineered Escherichia coli equipped with reconstructed one-carbon assimilation pathways. Proc. Natl Acad. Sci. USA. 115, E9271–E9279 (2018).

19. Bang, J., Hwang, C. H., Ahn, J. H., Lee, J. A. & Lee, S. Y. Escherichia coli is engineered to grow on CO2 and formic acid. Nat. Microbiol. 5, 1459–1463 (2020).

20. Kim, S. et al. Growth of E. coli on formate and methanol via the reductive glycine pathway. Nat. Chem. Biol. 16, 538–545 (2020).

21. Weinstock, M. T., Hesek, E. D., Wilson, C. M. & Gibson, D. G. Vibrio natriegens as a fast-growing host for molecular biology. Nat. Methods 13, 849–851, doi:10.1038/nmeth.3970 (2016).

22. Lee, H. H. et al. Functional genomics of the rapidly replicating bacterium Vibrio natriegens by CRISPRi. Nat. Microbiol. 4, 1105–1113 (2019).

23. Hoff, J. et al. Vibrio natriegens: an ultrafast-growing marine bacterium as emerging synthetic biology chassis. Environ. Microbiol. 22, 4394–4408 (2020).

24. Thoma, F. & Blombach, B. Metabolic engineering of Vibrio natriegens. Essays Biochem. 65, 381–392 (2021).

25. Xu, J. et al. Vibrio natriegens as a pET-compatible expression host complementary to Escherichia coli. Front. Microbiol. 12, 627181 (2021).

26. Eagon, R. G. & Wang, C. H. Dissimilation of glucose and gluconic acid by Pseudomonas natriegens. J. Bacteriol. 83, 879–886, doi:10.1128/jb.83.4.879-886.1962 (1962).

27. Hoffart, E. et al. High substrate uptake rates empower Vibrio natriegens as production host for industrial biotechnology. Appl. Environ. Microbiol. 83, doi:10.1128/aem.01614-17 (2017).

28. Dalia, T. N. et al. Multiplex genome editing by natural transformation (MuGENT) for synthetic biology in Vibrio natriegens. ACS Synth. Biol. 6, 1650–1655 (2017).

29. Zhang, Y. et al. Systems metabolic engineering of Vibrio natriegens for the production of 1,3-propanediol. Metab. Eng. 65, 52–65 (2021).

30. Akram, M. Citric acid cycle and role of its intermediates in metabolism. Cell Biochem. Biophys. 68, 475–478 (2014).

31. Vuoristo, K. S., Mars, A. E., Sanders, J. P. M., Eggink, G. & Weusthuis, R. A. Metabolic engineering of TCA cycle for production of chemicals. Trends Biotechnol. 34, 191–197 (2016).

32. Kroll, R. G. & Booth, I. R. The relationship between intracellular pH, the pH gradient and potassium transport in Escherichia coli. Biochem. J. 216, 709–716 (1983).

33. Warnecke, T. & Gill, R. T. Organic acid toxicity, tolerance, and production in Escherichia coli biorefining applications. Microb. Cell Fact. 4, 25 (2005).

34. Xu, Y. et al. An acid-tolerance response system protecting exponentially growing Escherichia coli. Nat. Commun. 11, 1496 (2020).

35. Yishai, O., Bouzon, M., Döring, V. & Bar-Even, A. In vivo assimilation of one-carbon via a synthetic reductive glycine pathway in Escherichia coli. ACS Synth. Biol. 7, 2023–2028 (2018).

36. Jakes, K. S. The colicin E1 TolC box: identification of a domain required for colicin E1 cytotoxicity and TolC binding. J. Bacteriol. 199, e00412–16 (2017).

37. Li, Q. et al. A modified pCas/pTargetF system for CRISPR-Cas9-assisted genome editing in Escherichia coli. Acta Biochim. Biophys Sin (Shanghai) 53, 620–627 (2021).

38. Banerjee, D. et al. Genome-scale metabolic rewiring improves titers rates and yields of the non-native product indigoidine at scale. Nat. Commun. 11, 5385 (2020).

39. Hong, Y., Arbter, P., Wang, W., Rojas, L. N. & Zeng, A. P. Introduction of glycine synthase enables uptake of exogenous formate and strongly impacts the metabolism in Clostridium pasteurianum. Biotechnol. Bioeng. 118, 1366–1380 (2021).

40. Fink, C. et al. A shuttle-vector system allows heterologous gene expression in the thermophilic methanogen Methanothermobacter thermautotrophicus ΔH. mBio. 12, e0276621 (2021).

41. Yu, D., Xu, F., Valiente, J., Wang, S. & Zhan, J. An indigoidine biosynthetic gene cluster from Streptomyces chromofuscus ATCC 49982 contains an unusual IndB homologue. J. Ind. Microbiol. Biotechnol. 40, 159–168 (2013).

42. Wang, L. et al. Protein scaffold optimizes arrangement of constituent enzymes in indigoidine synthetic pathway to improve the pigment production. Appl. Microbiol. Biotechnol. 104, 10493–10502 (2020).

43. Xu, F., Gage, D. & Zhan, J. Efficient production of indigoidine in Escherichia coli. J. Ind. Microbiol. Biotechnol. 42, 1149–1155 (2015).

44. Wehrs, M. et al. Correction to: Production efficiency of the bacterial non-ribosomal peptide indigoidine relies on the respiratory metabolic state in S. cerevisiae. Microb. Cell Fact. 18, 218 (2019).

