## Supplementary Materials for "Discovery and remodeling of *Vibrio natriegens* as a microbial platform for efficient formic acid biorefinery"

Jinzhong Tian *et al.*

**The PDF file includes:**

Figs. S1 to S8

Tables S1 to S4

References

**Fig. S1.**

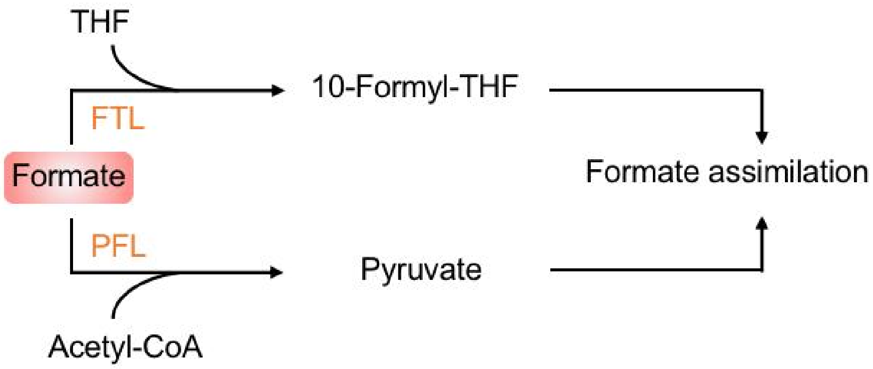


**Figure S1. The FTL and PFL-mediated formate assimilation reactions in microorganisms.**

FTL, formate tetrahydrofolate ligase. PFL, pyruvate formate-lyase.

**Fig. S2.**

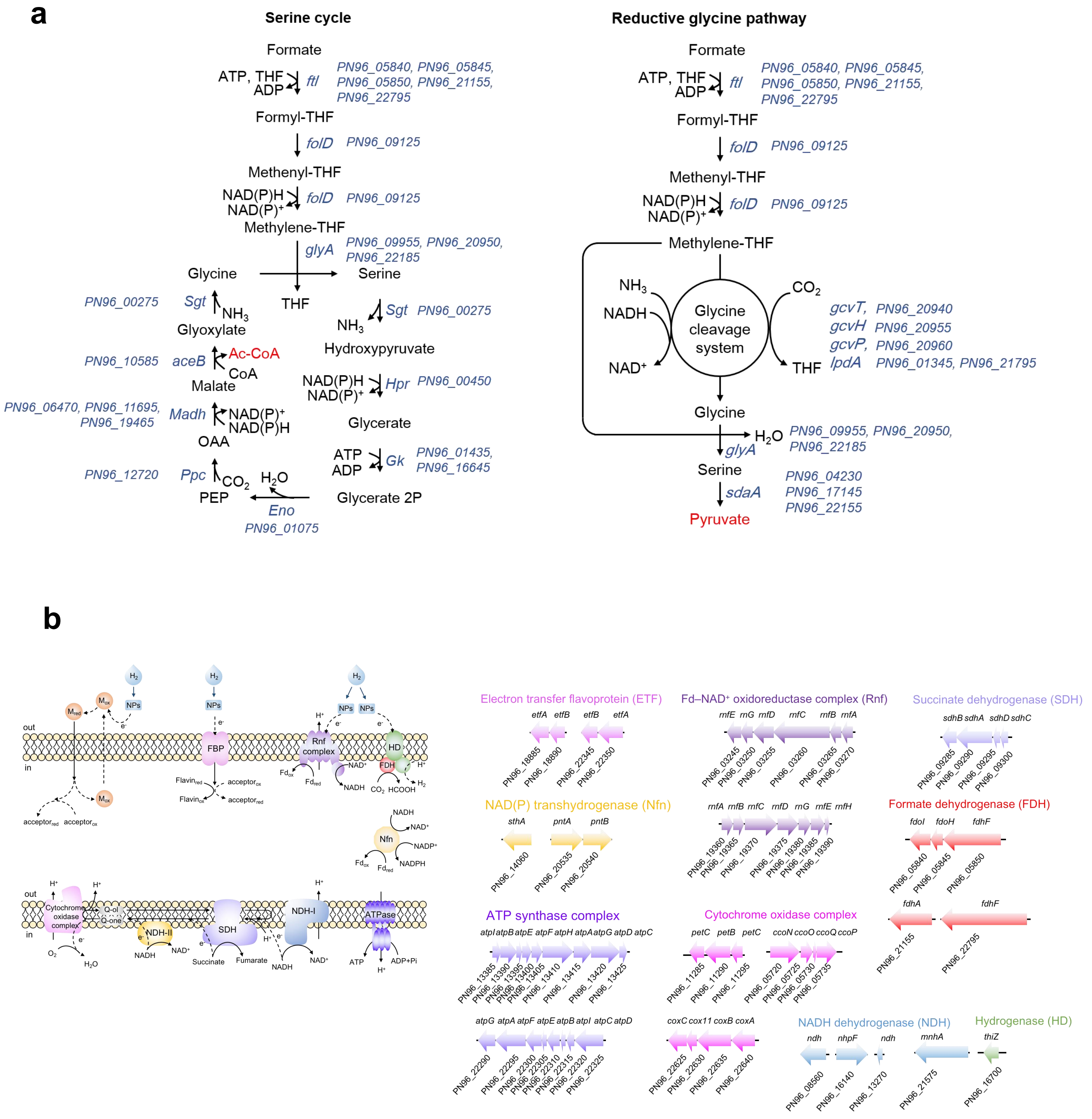


**Figure S2. The annotated genes associated with the serine cycle and reductive glycine pathway and energy conservation and supply in *V. natriegens*. a**, The genes responsible for serine cycle and reductive glycine pathway in *V. natriegens*. **b**, Arrangement of the genes associated with energy conservation and supply in *V. natriegens*. The putative energy conservation and supply systems in *V. natriegens* include metal ion mediated EET (extracellular electron transfer), flavin binding prtein (FBP), Fd-NAD+ oxidoreductase complex (Rnf), Hydrogenase (HD), formate dehydrogenase (FDH), transhydrogenase (Nfn), Cytochrome oxidase complex, NADH dehydrogenase II (NDH-II), succinate dehydrogenase (SDH), NADH dehydrogenase I (NDH-I), ATPase (ATP synthase), metal ion (M), and nano particles (NPs).

**Fig. S3.**

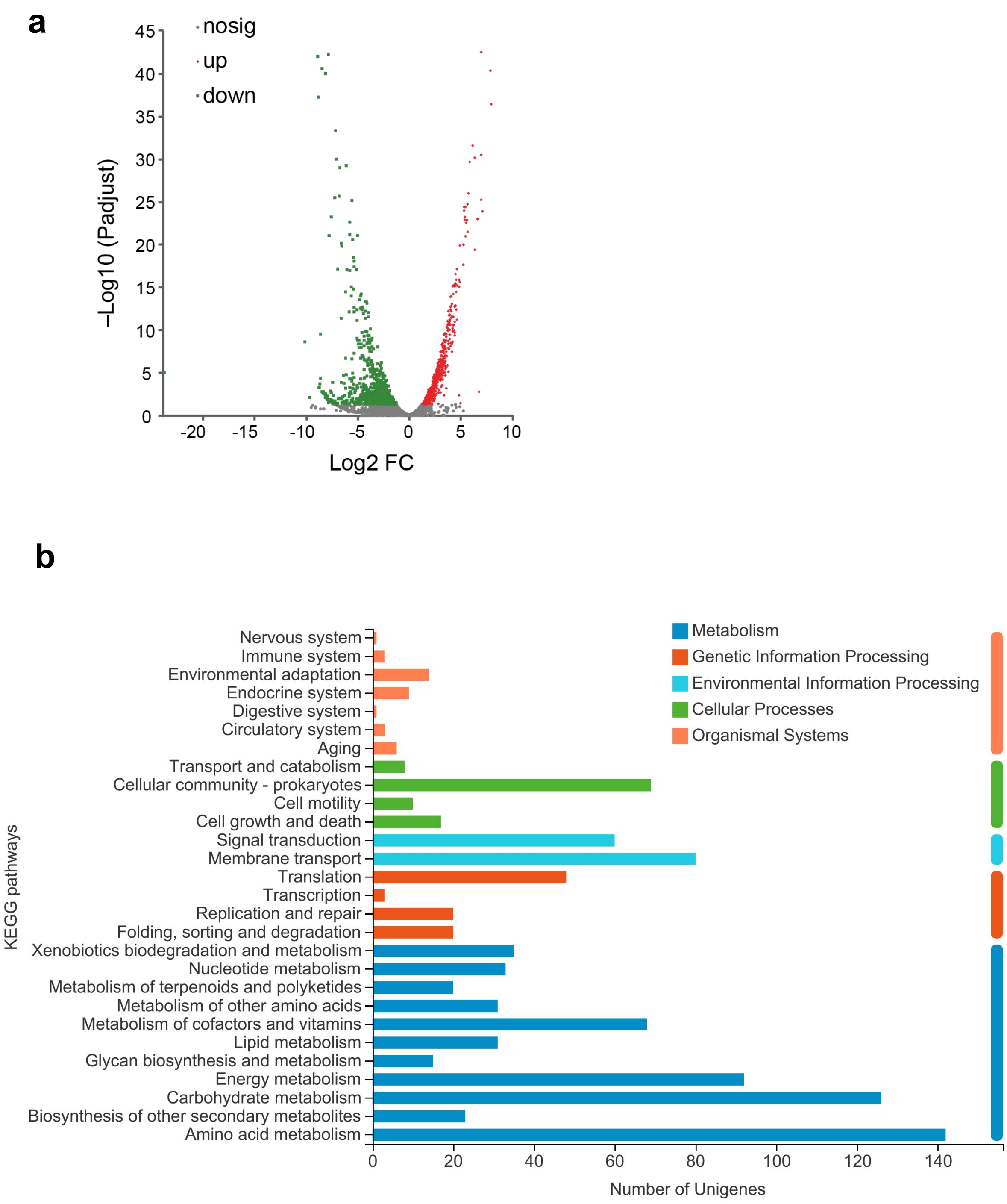


**Figure S3. Comparative transcriptomic analysis of *V. natriegens* in the presence and absence of formate using RNA-seq. a**, Volcano map of differentially expressed genes. Each dot represents one gene. The red and green dots indicate the genes exhibiting significantly upregulation and downregulation (FDR ≤ 0.05 and $|log_2FC| \geq 1$), respectively, with the supplementation of 40 $g \cdot L^{-1}$

sodium formate ($HCOONa \cdot 2H_2O$). The other genes were represented by grey dots. **b**, The top 28 functional enrichment subsets.

**Fig. S4.**

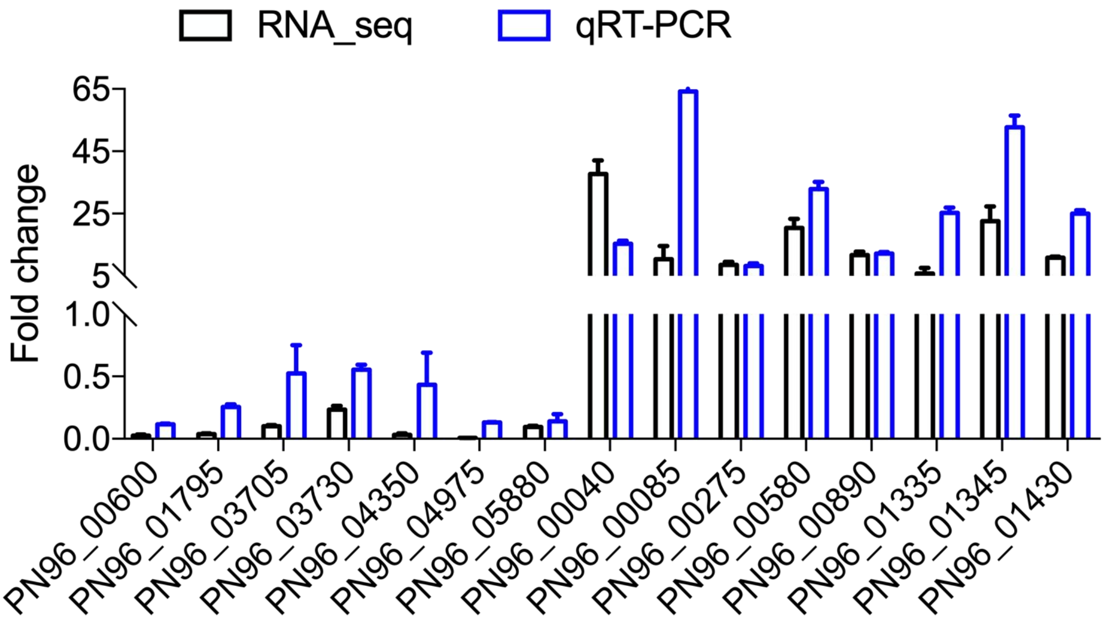


**Figure S4. Correlation between the RNA-seq and qRT-PCR results.** The *V. natriegens* strains were cultured in LBv2 medium containing 40 g $L^{-1}$ sodium formate ($HCOONa \cdot 2H_2O$). Cells were harvested after 6 h of fermentation. 15 genes that showed significantly changed transcriptional levels with formate stress (RNA-seq results) were picked out for the test. The data were mean ± standard deviation (SD) of 2 independent biological replicates.

**Fig. S5.**

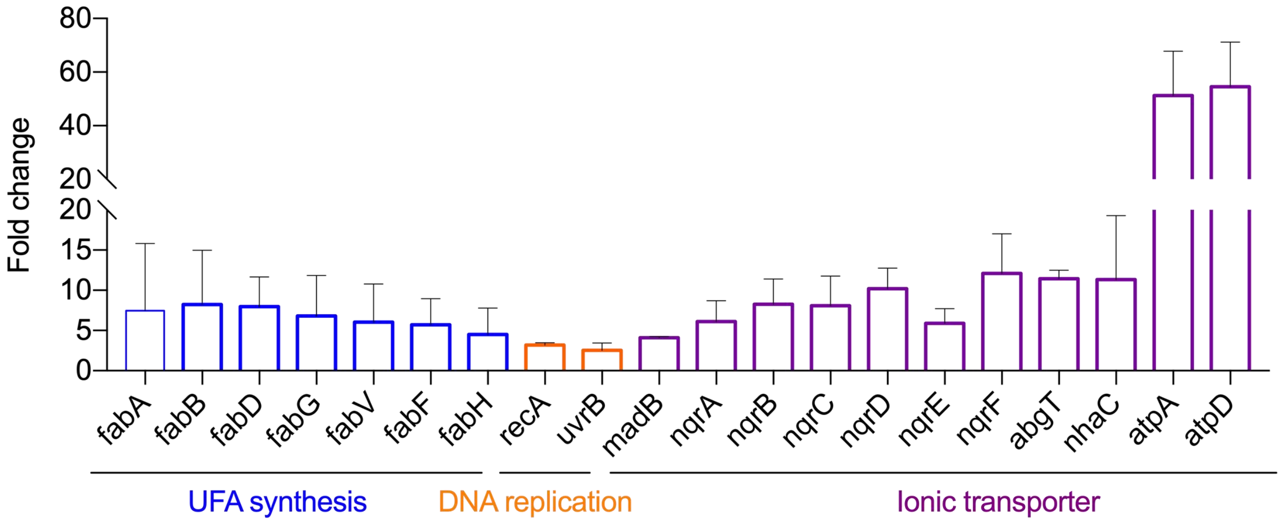


**Figure S5. Transcriptional differences of the *V. natriegens* genes responsible for ionic efflux systems, synthesis of unsaturated fatty acids and DNA repairing with and without the supplementation of sodium formate.** The data were mean ± standard deviation (SD) of 2 independent biological replicates from RNA-seq analysis.

### Fig. S6.

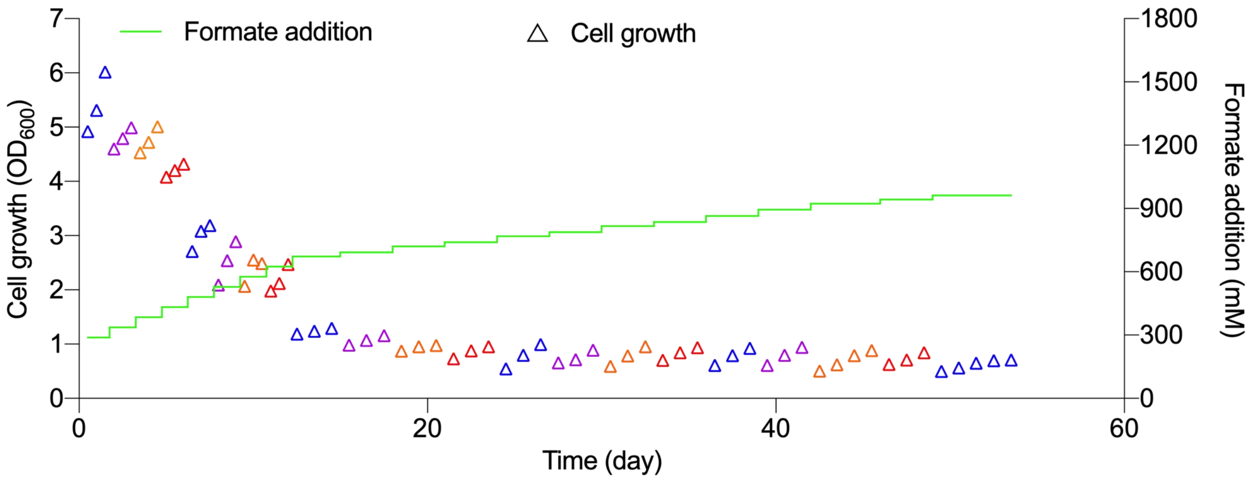


**Figure S6. The evolution trajectory of the wild-type *V. natriegens* strain for obtaining an evolved strain with enhanced formate utilization.** The LBv2 medium was used with an increasing concentration of sodium formate ($HCOONa \cdot 2H_2O$) from 30 to 100 $g \cdot L^{-1}$ (the equivalent of 288 to 962 mM formate). Triangles indicate the bacterial biomass of each passage after 12 h of cultivation, in which same colors represent the passages with the same sodium formate concentration.

**Fig. S7.**

a

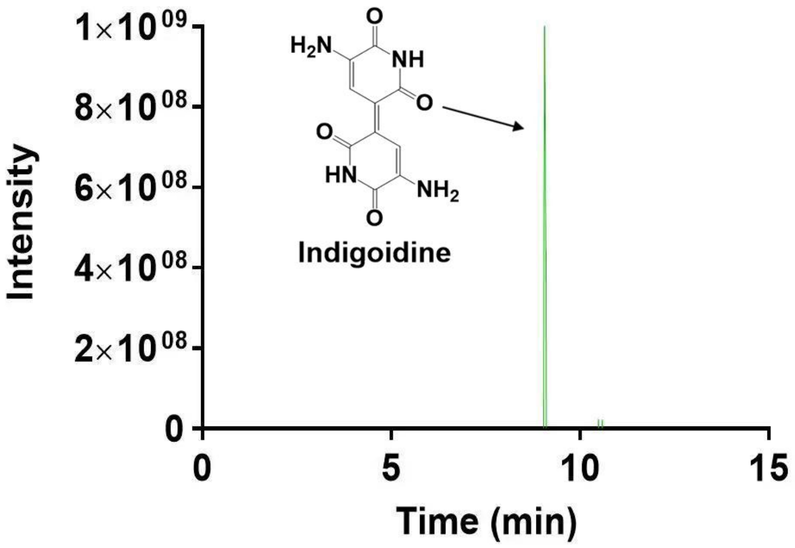


b

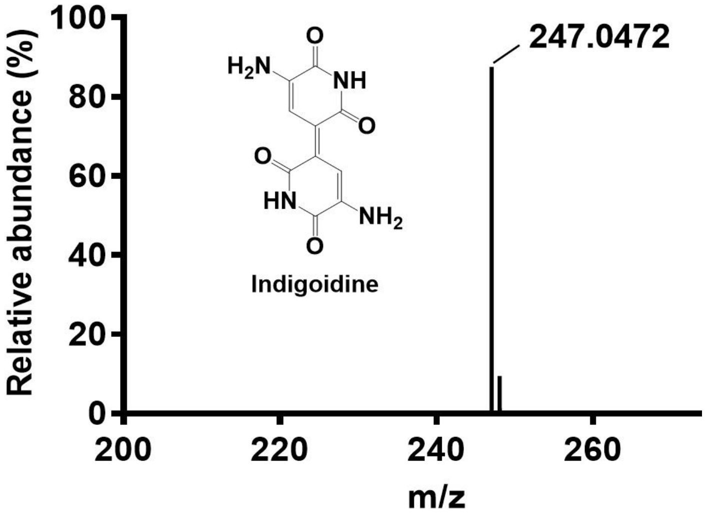


**Figure S7. High-performance liquid chromatography-mass spectrometry identification of the extracted indigoidine. a**, Detection of the extracted indigoidine by HPLC. **b**, Mass spectrometry analysis of the HPLC fraction corresponding to indigoidine.

**Fig. S8.**

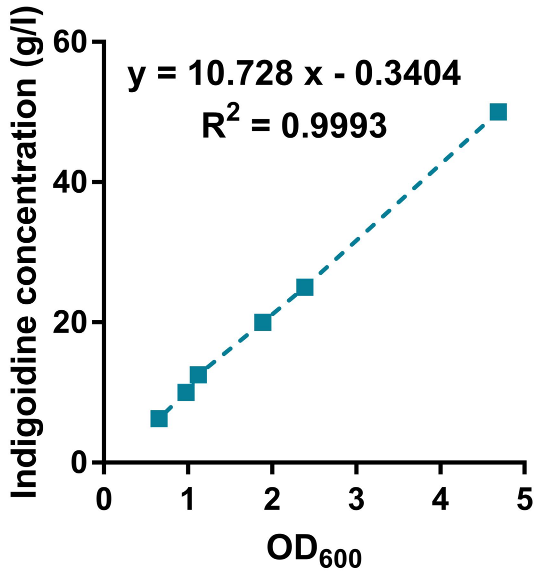


**Figure S8. Spectrophotometer standard curve correlating indigoidine concentration to the absorbance at 600 nm.**

**Table S1. Comparison of the tolerance and consumption of formate by the *V. natriegens* strains in the this study and other formate-utilizing microorganisms**

| Reported formate-utilizing microorganisms | Formate tolerance (mM) | Formate consumption ($mM \cdot h^{-1}$) | Carbon sources | References |
|---|---|---|---|---|
| *Vibrio natriegens* | 769.2 | 0.012 | Formate, yeast extract, peptone | This study |
| *Vibrio natriegens* -S-TCA-2.0-IE | 1480.8 | 71.1 | Formate, yeast extract, peptone | This study |
| *Vibrio natriegens* -S-TCA-2.1 | 712.3 | 11.1 | Formate | This study |
| *Saccharomyces cerevisiae* | 750 | 0 | Formate, glucose, $CO_2$ | (*12*) |
| *Umbelopsis isabellina* | 58.8* | 1.23 | Formate, yeast extract, glucose | (*11*) |
| *Thermococcus onnurineus* | 73.5* | ND | Formate | (*7*) |
| *Thermoacidophilic crenarchaeote* | 0.070* | 0.00017 | Formate | (*10*) |
| *Rhodopseudomonas palustris* | 0.01* | 0.0021 | Formate | (*5*) |
| *Ralstonia eutropha* | 43.5 | 0.95 | Formate | (*9*) |
| *Pseudomonas C* | 109* | ND | Formate | (*6*) |
| *Kuenenia stuttgartiensis* | 50* | ND | Formate, acetate, $CO_2$ | (*17*) |
| *Desulfurococcus amylolyticus* | 116.6* | ND | Formate | (*14*) |
| *Desulfovibrio desulfuricans* | 75* | ND | Formate, $CO_2$ | (*15*) |
| *Clostridium pasteurianum* | 117.6 | 0 | Formate, yeast extract, glucose | (*39*) |
| *Methanothermobacter spp.* | 200* | 3.95 | Formate | (*40*) |
| *Dehalococcoides mccartyi* | 2* | ND | Formate, acetate | (*8*) |
| *Acetobacterium wieringae* | 1* | ND | Formate, CO | (*13*) |

| | | | | |
|---|---|---|---|---|
| *Escherichia coli* | 0.0867* | 4.04 | Formate, $CO_2$, glucose | (*18*) |
| *Escherichia coli* | 0.0867* | 1.76 | Formate, $CO_2$ | (*19*) |
| *Escherichia coli* | 153 | 0.357 | Formate, $CO_2$ | (*20*) |

The artificial bacteria were highlighted with grey.

*Working concentration (the maximum tolerated concentration has not been tested).

**Table S2. Microbial synthesis of indigoidine reported previously**

| **Microorganisms** | Titer ($g \cdot L^{-1}$) | Productivity ($g \cdot L^{-1} \cdot h^{-1}$) | Carbon sources | Fermentation mode | References |
|---|---|---|---|---|---|
| *Escherichia coli* | 1.73 | 0.133 | Yeast extract, peptone | Batch | (*41*) |
| *Streptomyces coelicolor* | 0.59 | 0.004 | Sucrose, glucose, yeast extract | Batch | (*41*) |
| *Escherichia coli* | 14 | 0.233 | Yeast extract, glutamate | Batch | (*42*) |
| *Pseudomonas putida* | 25.6 | 0.22 | Glucose | Fed-batch | (*38*) |
| *Escherichia coli* | 8.81 | 0.315 | Yeast extract, peptone, glutamine | Batch | (*43*) |
| *Saccharomyces cerevisiae* | 0.98 | 0.014 | Yeast extract, peptone, glucose | Fed-batch | (*44*) |

**Table S3. Strains and plasmids used in this work**

| Strains/plasmids | Description/Genotype | Sources |
|---|---|---|
| **Strains** | | |
| *E. coli* DH5α | For plasmid construction | Invitrogen |
| *V. natriegens* ATCC 14048 | The parental *V. natriegens* strain | (*21*) |
| S-TCA-1.0 | Derived from *V. natriegens* ATCC 14048, *ΔPN96_00930ΔPN96_07295ΔPN96_10585ΔPN96_14755ΔPN96_06470ΔPN96_19465ΔPN96_11695::Km*$^{R}$*Δdns::Cm*$^{R}$ | This study |
| S-TCA-2.0 | Derived from S-TCA-1.0 by adaptive laboratory evolution | This study |
| S-TCA-2.1 | Derived from S-TCA-1.0 by adaptive laboratory evolution | This study |
| *V. natriegens Δfdh* | Derived from *V. natriegens* ATCC 14048, with the simultaneous deletion of three *fdh* genes (PN96_05850, PN96_21155, and PN96_22795) | |
| *V. natriegens Δftl* | Derived from *V. natriegens* ATCC 14048, with the deletion of the *ftl* gene (PN96_20840) | |
| *V. natriegens Δpfl* | Derived from *V. natriegens* ATCC 14048, with the deletion of the *pfl* gene (PN96_08455) | |
| *V. natriegens ΔfdhΔftlΔpfl* | Derived from *V. natriegens* ATCC 14048, with the simultaneous deletion of the abovementioned *ftl, pfl,* and the three *fdh* genes | |
| **Plasmids** | | |
| pColE1-Amp | Overexpression plasmid with pColE1 origin, ampicillin/carbenicillin resistance | (*36*) |
| pColE1-idgs-sfp | Derived from pColE1-Amp, Overexpression of the J23102-B00320m-idgs-RBS-sfp fragment | This study |
| pMMB67EH-tfox | The IPTG-inducible plasmid for expressing *tfoX*, ampicillin/carbenicillin resistance, pMMB origin | (*28*) |
| pMMB67EH-tfox-SacB | Derived from pMMB67EH-tfox, for *SacB* overexpression; | This study |

**Table S4. The primers used in this work**

| Names | Sequences (5'-3') | Description |
|---|---|---|
| Cm_F | GAAAGATTAGCGATTGTCGCGATTGGTGAGGATTAAAATACCTGTGACGGAAGATCAC | Forward primer for cloning the chloramphenicol resistance gene |
| Cm_R | TTTAAAGACTTTAACTATGAAATACCTGTTCTCTGTCGAATTTGCTTTCGAATTTCTGC | Reverse primer for cloning the chloramphenicol resistance gene |
| Kan_F | ATTGTCGCGATTGGTGAGGATTATTTGTTATCATTCTATAGTATTAAGTATTGTTCAGC | Forward primer for cloning the kanamycin resistance gene |
| Kan_R | CTTTTTAAAGACTTTAACTATGAAATACCTGTTCTCTAGTTCCTGCCCTCTGATTTTCC | Reverse primer for cloning the kanamycin resistance gene |
| KO_dns_UP_F | TAATCCTCACCAATCGCGAC | Forward primer for cloning the upstream homologous arm of the *dns* gene |
| KO_dns_UP_R | ACTGGTAAGCCATAACGACC | Reverse primer for cloning the upstream homologous arm of the *dns* gene |
| KO_dns_DN_F | CTAACATGGCTAAGCACCTG | Forward primer for cloning the downstream homologous arm of the *dns* gene |
| KO_dns_DN_R | AGAGAACAGGTATTTCATAGTTAAAGTC | Reverse primer for cloning the downstream homologous arm of the *dns* gene |
| KO_00930_DN_F | GGAAATTCCCTGCCTTTATTGTTTCTGTGCTTCTTTTTCTCGTCCTTAAAATGGAAAGG | Forward primer for cloning the downstream homologous arm of the *PN96_00930* gene |
| KO_00930_DN_R | GAACAATCTGTAATTAGTGGATCGTTG | Reverse primer for cloning the downstream homologous arm of the *PN96_00930* gene |
| KO_00930_UP_F | CACTAGAAAAAGATCCACGCATCC | Forward primer for cloning the upstream homologous arm of the *PN96_00930* gene |
| KO_00930_UP_R | GAAGCACAGAAACAATAAAGGCAG | Reverse primer for cloning the upstream homologous arm of the *PN96_00930* gene |
| KO_00930_seq_F | GGTGAGTAATCAGGCTGAGTATC | Forward primer for identifying the *PN96_00930* knockout |
| KO_00930_seq_R | CTAATGGTCTGTTTGCTCATCGTG | Reverse primer for identifying the *PN96_00930* knockout |

| | | |
|---|---|---|
| KO_14755_DN_F | AAAATAAAAAGAGAAACGCCCACGACAAAGATTTAAATGACATAAAAAAATGGGACGCG | Forward primer for cloning the downstream homologous arm of the *PN96_14755* gene |
| KO_14755_DN_R | CTACGGCAGCAATCAGAGAGTAAC | Reverse primer for cloning the downstream homologous arm of the *PN96_14755* gene |
| KO_14755_UP_F | ACTCAAGTGAGATGGCGTTTAAG | Forward primer for cloning the upstream homologous arm of the *PN96_14755* gene |
| KO_14755_UP_R | GTGGGCGTTTCTCTTTTTATTTTTAG | Reverse primer for cloning the upstream homologous arm of the *PN96_14755* gene |
| KO_14755_seq_F | CCGATCATCTCATTGAACGTATTC | Forward primer for identifying the *PN96_14755* knockout |
| KO_14755_seq_R | GTAAAGATCGCCCTGACGATATTC | Reverse primer for identifying the *PN96_14755* knockout |
| KO_07295_DN_F | TGGCGCGCATCCTTCAACCCTACATAAGGCTCCAAATTTAACCGCAAAAGAAAAACTCG | Forward primer for cloning the downstream homologous arm of the *PN96_07295* gene |
| KO_07295_DN_R | ACCATACAGTTTTCTTCGCCC | Reverse primer for cloning the downstream homologous arm of the *PN96_07295* gene |
| KO_07295_UP_F | CAAAGATGGGGGCGATGGTG | Forward primer for cloning the upstream homologous arm of the *PN96_07295* gene |
| KO_07295_UP_R | TTGGAGCCTTATGTAGGGTTGAAG | Reverse primer for cloning the upstream homologous arm of the *PN96_07295* gene |
| KO_07295_seq_F | GAGAATCACGCGACTTTGAAAC | Forward primer for identifying the *PN96_07295* knockout |
| KO_07295_seq_R | CTCATCATATCTGATACAAACAAAAAG | Reverse primer for identifying the *PN96_07295* knockout |
| KO_10585_DN_F | CAAGATTGATAAAGCACGTTAAGGATGAATGATTGAATACGAATACCAGAAAACAATAC | Forward primer for cloning the downstream homologous arm of the *PN96_10585* gene |
| KO_10585_DN_R | CAAAAAGCCAATCGCCGTC | Reverse primer for cloning the downstream homologous arm of the *PN96_10585* gene |

| | | |
|---|---|---|
| KO_10585_UP_F | CTCCAGAGCTTGCTCTGCGTTAC | Forward primer for cloning the upstream homologous arm of the *PN96_10585* gene |
| KO_10585_UP_R | AATCATTCATCCTTAACGTGCTTTATC | Reverse primer for cloning the upstream homologous arm of the *PN96_10585* gene |
| KO_10585_seq_R | GCTGTGCAATTGTGTTTGCAG | Forward primer for identifying the *PN96_10585* knockout |
| KO_10585_seq_F | CATCAAAAATCAACCGTTTGATCCG | Reverse primer for identifying the *PN96_10585* knockout |
| KO_06470_DN_F | CTTTCGACTACTTGTTATTTTAGAGAGAGATTACACCCCATCTCCTGCGGTTTTCAACG | Forward primer for cloning the downstream homologous arm of the *PN96_06470* gene |
| KO_06470_DN_R | ATTTGAACGGGTAATATAGTTTGCC | Reverse primer for cloning the downstream homologous arm of the *PN96_06470* gene |
| KO_06470_UP_F | ATTCACTAACCCCCGAAAAAGTAG | Forward primer for cloning the upstream homologous arm of the *PN96_06470* gene |
| KO_06470_UP_R | TGGGGTGTAATCTCTCTCTAAAATAAC | Reverse primer for cloning the upstream homologous arm of the *PN96_06470* gene |
| KO_06470_seq_F | GCTCTCTAACAAATGGGCTTTG | Forward primer for identifying the *PN96_06470* knockout |
| KO_06470_seq_R | CCACAAGAGCATTTGGGTTTG | Reverse primer for identifying the *PN96_06470* knockout |
| KO_19465_DN_F | CAACTACTTGTTATTTTAGAGAGATTACACCCCACGCCTTACGGTTTTGGTCGTTATAC | Forward primer for cloning the downstream homologous arm of the *PN96_19465* gene |
| KO_19465_DN_R | AAGTAAACAAAGCGGTATAGACATTAG | Reverse primer for cloning the downstream homologous arm of the *PN96_19465* gene |
| KO_19465_UP_F | TTTCTTGCCGTTTCATGGACAG | Forward primer for cloning the upstream homologous arm of the *PN96_19465* gene |
| KO_19465_UP_R | TGGGGTGTAATCTCTCTAAAATAACAAG | Reverse primer for cloning the upstream homologous arm of the *PN96_19465* gene |

| | | |
|---|---|---|
| KO_19465_seq_F | GCAATTTTAACAATAAGAACAAACACG | Forward primer for identifying the *PN96_19465* knockout |
| KO_19465_seq_R | CAAAAAAGTTATCGCGTCTAACG | Reverse primer for identifying the *PN96_19465* knockout |
| KO_11695_DN_F | GTTTAAGCGTTTATACGTTTATAAAAAGC | Forward primer for cloning the downstream homologous arm of the *PN96_11695* gene |
| KO_11695_DN_R | CAGCAGATACAGCGCGAATAAAG | Reverse primer for cloning the downstream homologous arm of the *PN96_11695* gene |
| KO_11695_UP_F | TAGATATAAGACGCTGAAAAGCTACC | Forward primer for cloning the upstream homologous arm of the *PN96_11695* gene |
| KO_11695_UP_R | GTAGTTCTCCTTGAGAGTATTTTTTATAAATG | Reverse primer for cloning the upstream homologous arm of the *PN96_11695* gene |
| KO_11695_kanR_F | AAAATACTCTCAAGGAGAACTACTTTGTTATCATTCTATAGTATTAAGTATTGTTCAGC | Forward primer for cloning the kanamycin resistance gene |
| KO_11695_kanR_R | TAAGATCGGCTTTTTATAAACGTATAAACGCTTAAACAGTTCCTGCCCTCTGATTTTCC | Reverse primer for cloning the kanamycin resistance gene |
| KO_11695_seq_F | GTGATTCCTATAGCTTTACTTGGTTG | Forward primer for identifying the *PN96_11695* knockout |
| KO_11695_seq_R | CGAAGCGATCGAGAAAGCAAAC | Reverse primer for identifying the *PN96_11695* knockout |
| KO_20840_DN_F | ACGGTTTAGTCACTTTGGGAGCATCCCATCACATCCTCCTGTCATTTAAATTAAACTTC | Forward primer for cloning the downstream homologous arm of the *PN96_20840* gene |
| KO_20840_DN_R | CGGAGCTTGCTCAGTGGAATG | Reverse primer for cloning the downstream homologous arm of the *PN96_20840* gene |
| KO_20840_UP_F | GATGTTCGATATATTGCGAGAGCC | Forward primer for cloning the upstream homologous arm of the *PN96_20840* gene |
| KO_20840_UP_R | GGATGCTCCCAAAGTGACTAAAC | Reverse primer for cloning the upstream homologous arm of the *PN96_20840* gene |
| KO_20840_seq_F | GCGAAGACAGTTTAGAAGAGGC | Forward primer for identifying the *PN96_20840* knockout |

| | | |
|---|---|---|
| KO_20840_seq_R | CAATGTAAAGTAATGCCGAAACAATG | Reverse primer for identifying the *PN96_20840* knockout |
| KO_08455_DN_R | GCATTACTTGGATCGAGTTGAGAC | Forward primer for cloning the downstream homologous arm of the *PN96_08455* gene |
| KO_08455_DN_F | TACGTATTTTTTTCTACTAAAAAGGTAGGTAT GTCGACTGTCGCGAAATAACGTTATAG | Reverse primer for cloning the downstream homologous arm of the *PN96_08455* gene |
| KO_08455_UP_R | GACATACCTACCTTTTTAGTAGAAAAAAATA CG | Forward primer for cloning the upstream homologous arm of the *PN96_08455* gene |
| KO_08455_UP_F | CGGATGGCGTAGATGACGAAC | Reverse primer for cloning the upstream homologous arm of thePN96_08455 gene |
| KO_08455_seq_F | CTTGTCGCTACCCCATTTTTATC | Forward primer for identifying the *PN96_08455* knockout |
| KO_08455_Seq_R | CGTTGGTGTAATCATGGTCAAC | Reverse primer for identifying the *PN96_08455* knockout |
| KO_05880_DN_F | GTGAGCAAGTAGCTCCTTCACTGAACAGGAC AAGACTTTTGCAAAAGCAAAGTTTGGTG | Forward primer for cloning the downstream homologous arm of the *PN96_05880* gene |
| KO_05880_DN_R | GTATCGAGGTTCGCGTGATTTATTC | Reverse primer for cloning the downstream homologous arm of the *PN96_05880* gene |
| KO_05880_UP_F | CAAATTTAAGAGACGGAGACTGACC | Forward primer for cloning the upstream homologous arm of the *PN96_05880* gene |
| KO_05880_UP_R | AGTCTTGTCCTGTTCAGTGAAGG | Reverse primer for cloning the upstream homologous arm of the *PN96_05880* gene |
| KO_05880_seq_F | GTATTAAGTAGCGTGAATCTCTCGC | Forward primer for identifying the *PN96_05880* knockout |
| KO_05880_seq_R | CCTTTGTGAATAAACGGTATTAGCG | Reverse primer for identifying the *PN96_05880* knockout |
| KO_05840_50_UP_F | GTTCTTGCGACTAAGCGCATTTTC | Forward primer for cloning the downstream homologous arm of the *PN96_05840*, *PN96_05845*, *PN96_05850* genes |

| | | |
|---|---|---|
| KO_05840_50_UP_R | CGGGATTTGGATTGCGTACATC | Reverse primer for cloning the downstream homologous arm of the *PN96_05840*, *PN96_05845*, *PN96_05850* genes |
| KO_05850_50_DN_F | GTAAAACAACTCAGGATGTACGCAATCCAAATCCCGCTCTTACCTCCTAAAGTGTGTCG | Forward primer for cloning the upstream homologous arm of the *PN96_05840*, *PN96_05845*, *PN96_05850* genes |
| KO_05850_50_DN_R | CAAGAAATCGTCGAACCTGCTC | Reverse primer for cloning the upstream homologous arm of the *PN96_05840*, *PN96_05845*, *PN96_05850* genes |
| KO_05850_50_Seq_F | CCAAACGATAACCAGCATAAATAATAGC | Forward primer for identifying the knockout of *PN96_05840*, *PN96_05845*, and *PN96_05850* |
| KO_05850_50_seq_R | CAGGAACTACGAAAGTGGCAAAC | Reverse primer for identifying the knock of *PN96_05840*, *PN96_05845*, and *PN96_05850* |
| KO_21155_DN_F | AGTTCATCTAAAGAATCTAACAGGTAAGTCTTTGATGATTGAGAAGTGGAATTGATGGC | Forward primer for cloning the downstream homologous arm of the *PN96_21155* gene |
| KO_21155_DN_R | GATTACATAGATAGCAGACTGCTTTTTATTTTAC | Reverse primer for cloning the downstream homologous arm of the *PN96_21155* gene |
| KO_21155_UP_F | CACCGATCCGGACTGGGTGC | Forward primer for cloning the upstream homologous arm of the *PN96_21155* gene |
| KO_21155_UP_R | CAAAGACTTACCTGTTAGATTCTTTAG | Reverse primer for cloning the upstream homologous arm of the *PN96_21155* gene |
| KO_21155_Seq_F | GAAGAACCCCTTAGCCGAATG | Forward primer for identifying the *PN96_21155* knockout |
| KO_21155_Seq_R | CAACGGGTAACTGTGGACTG | Reverse primer for identifying the *PN96_21155* knockout |
| KO_22795_DN_F | TTCGAGGTTCAATGCGAAGCCCTTTTTATCCTGTCTGCACGCTACCCTGAGTTAGCCAC | Forward primer for cloning the downstream homologous arm of the *PN96_22795* gene |
| KO_22795_DN_R | CGTAATCAACCGTACCTTGCC | Reverse primer for cloning the |

| | | |
|---|---|---|
| | | downstream homologous arm of the *PN96_22795* gene |
| KO_22795_UP_F | CAGCGAAAGCAGATGTTGCG | Forward primer for cloning the upstream homologous arm of the *PN96_22795* gene |
| KO_22795_UP_R | GCAGACAGGATAAAAAGGGC | Reverse primer for cloning the upstream homologous arm of the *PN96_22795* gene |
| KO_22795_seq_F | CTGAATAACAGAGGGTCGTCAG | Forward primer for identifying the *PN96_22795* knockout |
| KO_22795_seq_R | GTTAGTGAGCATTTTGTCACATGAAC | Reverse primer for identifying the *PN96_22795* knockout |
| tfox_F | AAAGAAAATGCCGATACTCGAGCTTCTGCTCCCGA | Forward primer for cloning the DNA fragment of *tfox* |
| tfox_R | TCTGGCCTATGGAGCTGTGCGGCAGCGCTCAGTAGG | Reverse primer for cloning the DNA fragment of *tfox* |
| sacB_F | GCTCCATAGGCCAGATCTTTAGGCCCGTAGTCTGCAA | Forward primer for cloning the DNA fragment of *sacB* |
| sacB_R | GGAGCAGAAGCTCGAGTATCGGCATTTTCTTTTGCGT | Reverse primer for cloning the DNA fragment of *sacB* |
| pColE1-AmpR-F | TCTAGAGGATCCCCGGGTAC | Forward primer for cloning the linear plasmid pColE1-Amp vector |
| pColE1-AmpR-R | GAGGATATCACAATGGTGTCGTACGAAGAGCTTTTATAGCATGTGAGCAAAAGGCCAGC | Reverse primer for cloning the linear plasmid pColE1-Amp vector |
| idgs-sfp-F | ATTTTGTAGAGTCACACAGGAAAGTACTAATACCATGATGACCCTTCAAGAAACCAGCG | Forward primer for cloning the DNA fragment of idgs-sfp |
| idgs-sfp-R | CTATAAAAGCTCTTCGTACGACACC | Reverse primer for cloning the DNA fragment of idgs-sfp |
| J23102-B0032m-F | GAATTCGAGCTCGGTACCCGGGGATCCTCTAGATTGACAGCTAGCTCAGTCCTAGGTACTGTGC | Forward primer for cloning the DNA fragment of J23102-B0032m |
| J23102-B0032m-R | TTAGTACTTTCCTGTGTGACTCTACAAAATTATTGCTAGCACAGTACCTAGGACTGAGC | Reverse primer for cloning the DNA fragment of J23102-B0032m |
| RT00600-F | ACTGGCGACATCTTGATTGGT | Forward primer for *PN96_00600* RT-PCR |
| RT00600-R | AAAGCGGGAACACTTGTCCA | Reverse primer for *PN96_00600* RT-PCR |
| RT04975-F | TATCCCGGTAGAGGCTCTGG | Forward primer for *PN96_04975* |

| | | |
|---|---|---|
| | | RT-PCR |
| RT04975-R | CCCTGCCGTTTCTACCACAT | Reverse primer for *PN96_04975* RT-PCR |
| RT05880-F | CCCATTCTGCCGCAGTGATA | Forward primer for *PN96_05880* RT-PCR |
| RT05880-R | GACTGCTTAAGCGGCACTTG | Reverse primer for *PN96_05880* RT-PCR |
| RT00275-F | AAACGTTGTTCGTGCTGGTG | Forward primer for *PN96_00275* RT-PCR |
| RT00275-R | TGACGTAACTGCGTCGACAA | Reverse primer for *PN96_00275* RT-PCR |
| RT01795-F | AGGGGCAATGTTGACCTACG | Forward primer for *PN96_01795* RT-PCR |
| RT01795-R | CGTTCAGTAATGTGCGCGAG | Reverse primer for *PN96_01795* RT-PCR |
| RT00890-F | CTGATCGCGTTCTGGGATGA | Forward primer for *PN96_00890* RT-PCR |
| RT00890-R | CACCGTGACAGTCGTGAGAA | Reverse primer for *PN96_00890* RT-PCR |
| RT01355-F | TTCAAAGGTGCTGGCTGGAA | Forward primer for *PN96_01355* RT-PCR |
| RT01355-R | ACCACGCTTAAGTGCGAAGA | Reverse primer for *PN96_01355* RT-PCR |
| RT01430-F | TCGAAACTGAACATGGGCGA | Forward primer for *PN96_01430* RT-PCR |
| RT01430-R | CAGCAACGTCACCTGGTTTG | Reverse primer for *PN96_01430* RT-PCR |
| RT04350-F | TGACGTTATCGCGTTCACCA | Forward primer for *PN96_04350* RT-PCR |
| RT04350-R | CCATCTCACCAAGCGCCTTA | Reverse primer for *PN96_04350* RT-PCR |
| RT03705-F | TGAGCAGCACCCTAGAAACG | Forward primer for *PN96_03705* RT-PCR |
| RT03705-R | ACTCAATTCACCGCTCTGGG | Reverse primer for *PN96_03705* RT-PCR |
| RT03730-F | TGCCAAGGGATACCGACAAC | Forward primer for *PN96_03730* RT-PCR |
| RT03730-R | GGCCCAGTTGTGAGGCTTTA | Reverse primer for *PN96_03730* |

| | | |
|---|---|---|
| | | RT-PCR |
| RT00040-F | ACAACGCGGTTTGATTGGTG | Forward primer for *PN96_00040* RT-PCR |
| RT00040-R | AATTGCGCCATTGAGTCTGC | Reverse primer for *PN96_00040* RT-PCR |
| RT00085-F | CTGCAACAGATGGCCCAATG | Forward primer for *PN96_00085* RT-PCR |
| RT00085-R | GCGCTTCTGCTAGTTCAACG | Reverse primer for *PN96_00085* RT-PCR |
| RT00580-F | TTCGCTCGTACTTTACGCGA | Forward primer for *PN96_00580* RT-PCR |
| RT00580-R | CTAGAGGGCGACGCATCTTC | Reverse primer for *PN96_00580* RT-PCR |
| RT00205-R | GTTAGCGACAGAAGAAGCAC | Forward primer for *PN96_00205* RT-PCR |
| RT00205-R | CCGGTATTCCTTCAGATCTC | Reverse primer for *PN96_00205* RT-PCR |
| RT01345-R | ATACGTGGGTCTTCATGCGG | Reverse primer for *PN96_01345* RT-PCR |

**References:**


1. Jiang, W. et al. Metabolic engineering strategies to enable microbial utilization of C1 feedstocks. *Nat. Chem. Biol.* **17**, 845-855 (2021).
2. Yishai, O., Goldbach, L., Tenenboim, H., Lindner, S. N. & Bar-Even, A. Engineered assimilation of exogenous and endogenous formate in *Escherichia coli*. *ACS Synth. Biol.* **6**, 1722-1731 (2017).
3. Bang, J. et al. Synthetic formatotrophs for one-carbon biorefinery. *Adv. Sci. (Weinh)* **8**, 2100199 (2021).
4. Bar-Even, A. Formate assimilation: the metabolic architecture of natural and synthetic pathways. *Biochemistry* **55**, 3851-3863 (2016).
5. Stokes, J. E. & Hoare, D. S. Reductive pentose cycle and formate assimilation in *Rhodopseudomonas palustris*. *J. Bacteriol.* **100**, 890-894 (1969).
6. Goldberg, I. & Mateles, R. I. Growth of *Pseudomonas C* on C1 compounds: enzyme activites in extracts of *Pseudomonas C* cells grown on methanol, formaldehyde, and formate as sole carbon sources. *J. Bacteriol.* **122**, 47-53 (1975).
7. Moon, Y. J. et al. Proteome analyses of hydrogen-producing hyperthermophilic archaeon *Thermococcus onnurineus* NA1 in different one-carbon substrate culture conditions. *Mol. Cell. Proteomics* **11**, M111.015420 (2012).
8. Zhuang, W. Q. et al. Incomplete Wood-Ljungdahl pathway facilitates one-carbon metabolism in organohalide-respiring *Dehalococcoides mccartyi*. *Proc. Natl Acad. Sci. USA* **111**, 6419-6424 (2014).
9. Grunwald, S. et al. Kinetic and stoichiometric characterization of organoautotrophic growth of *Ralstonia eutropha* on formic acid in fed-batch and continuous cultures. *Microb. Biotechnol.* **8**, 155-163 (2015).
10. Urschel, M. R., Hamilton, T. L., Roden, E. E. & Boyd, E. S. Substrate preference, uptake kinetics and bioenergetics in a facultatively autotrophic, *thermoacidophilic crenarchaeote*. *FEMS. Microbiol. Ecol.* **92**, fiw069 (2016).
11. Liu, Z. et al. Exploring eukaryotic formate metabolisms to enhance microbial growth and lipid accumulation. *Biotechnol. Biofuels* **10**, 22 (2017).
12. Gonzalez de la Cruz, J., Machens, F., Messerschmidt, K. & Bar-Even, A. Core catalysis of the reductive glycine pathway demonstrated in Yeast. *ACS Synth. Biol.* **8**, 911-917 (2019).
13. Arantes, A. L. et al. Enrichment of anaerobic syngas-converting communities and isolation of a novel carboxydotrophic *Acetobacterium wieringae* strain JM. *Front. Microbiol.* **11**, 58

(2020).

14. Ergal, I. et al. Formate utilization by the crenarchaeon *Desulfurococcus amylolyticus*. *Microorganisms* **8**, 454 (2020).

15. Sánchez-Andrea, I. et al. The reductive glycine pathway allows autotrophic growth of *Desulfovibrio desulfuricans*. *Nat. Commun.* **11**, 5090 (2020).

16. Wood, G. E., Haydock, A. K. & Leigh, J. A. Function and regulation of the formate dehydrogenase genes of the methanogenic archaeon *Methanococcus maripaludis*. *J. Bacteriol.* **185**, 2548-2554 (2003).

17. Lawson, C. E. et al. Autotrophic and mixotrophic metabolism of an anammox bacterium revealed by in vivo $^{13}C$ and $^{2}H$ metabolic network mapping. *Isme j.* **15**, 673-687 (2021).

18. Bang, J. & Lee, S. Y. Assimilation of formic acid and $CO_2$ by engineered *Escherichia coli* equipped with reconstructed one-carbon assimilation pathways. *Proc. Natl Acad. Sci. USA.* **115**, E9271-E9279 (2018).

19. Bang, J., Hwang, C. H., Ahn, J. H., Lee, J. A. & Lee, S. Y. *Escherichia coli* is engineered to grow on $CO_2$ and formic acid. *Nat. Microbiol.* **5**, 1459-1463 (2020).

20. Kim, S. et al. Growth of *E. coli* on formate and methanol via the reductive glycine pathway. *Nat. Chem. Biol.* **16**, 538-545 (2020).

21. Weinstock, M. T., Hesek, E. D., Wilson, C. M. & Gibson, D. G. *Vibrio natriegens* as a fast-growing host for molecular biology. *Nat. Methods* **13**, 849-851, doi:10.1038/nmeth.3970 (2016).

22. Lee, H. H. et al. Functional genomics of the rapidly replicating bacterium *Vibrio natriegens* by CRISPRi. *Nat. Microbiol.* **4**, 1105-1113 (2019).

23. Hoff, J. et al. *Vibrio natriegens*: an ultrafast-growing marine bacterium as emerging synthetic biology chassis. *Environ. Microbiol.* **22**, 4394-4408 (2020).

24. Thoma, F. & Blombach, B. Metabolic engineering of *Vibrio natriegens*. *Essays Biochem.* **65**, 381-392 (2021).

25. Xu, J. et al. *Vibrio natriegens* as a pET-compatible expression host complementary to *Escherichia coli*. *Front. Microbiol.* **12**, 627181 (2021).

26. Eagon, R. G. & Wang, C. H. Dissimilation of glucose and gluconic acid by *Pseudomonas natriegens*. *J. Bacteriol.* **83**, 879-886, doi:10.1128/jb.83.4.879-886.1962 (1962).

27. Hoffart, E. et al. High substrate uptake rates empower *Vibrio natriegens* as production host for industrial biotechnology. *Appl. Environ. Microbiol.* **83**, doi:10.1128/aem.01614-17 (2017).

28. Dalia, T. N. et al. Multiplex genome editing by natural transformation (MuGENT) for synthetic biology in *Vibrio natriegens*. *ACS Synth. Biol.* **6**, 1650-1655 (2017).
29. Zhang, Y. et al. Systems metabolic engineering of *Vibrio natriegens* for the production of 1,3-propanediol. *Metab. Eng.* **65**, 52-65 (2021).
30. Akram, M. Citric acid cycle and role of its intermediates in metabolism. *Cell Biochem. Biophys.* **68**, 475-478 (2014).
31. Vuoristo, K. S., Mars, A. E., Sanders, J. P. M., Eggink, G. & Weusthuis, R. A. Metabolic engineering of TCA cycle for production of chemicals. *Trends Biotechnol.* **34**, 191-197 (2016).
32. Kroll, R. G. & Booth, I. R. The relationship between intracellular pH, the pH gradient and potassium transport in *Escherichia coli. Biochem. J.* **216**, 709-716 (1983).
33. Warnecke, T. & Gill, R. T. Organic acid toxicity, tolerance, and production in *Escherichia coli* biorefining applications. *Microb. Cell Fact.* **4**, 25 (2005).
34. Xu, Y. et al. An acid-tolerance response system protecting exponentially growing *Escherichia coli. Nat. Commun.* **11**, 1496 (2020).
35. Yishai, O., Bouzon, M., Döring, V. & Bar-Even, A. In vivo assimilation of one-carbon via a synthetic reductive glycine pathway in *Escherichia coli*. *ACS Synth. Biol.* **7**, 2023-2028 (2018).
36. Jakes, K. S. The colicin E1 TolC box: identification of a domain required for colicin E1 cytotoxicity and TolC binding. *J. Bacteriol.* **199**, e00412-16 (2017).
37. Li, Q. et al. A modified pCas/pTargetF system for CRISPR-Cas9-assisted genome editing in *Escherichia coli. Acta Biochim. Biophys Sin (Shanghai)* **53**, 620-627 (2021).
38. Banerjee, D. et al. Genome-scale metabolic rewiring improves titers rates and yields of the non-native product indigoidine at scale. *Nat. Commun.* **11**, 5385 (2020).
39. Hong, Y., Arbter, P., Wang, W., Rojas, L. N. & Zeng, A. P. Introduction of glycine synthase enables uptake of exogenous formate and strongly impacts the metabolism in *Clostridium pasteurianum*. *Biotechnol. Bioeng.* **118**, 1366-1380 (2021).
40. Fink, C. et al. A shuttle-vector system allows heterologous gene expression in the thermophilic methanogen *Methanothermobacter thermautotrophicus* ΔH. *mBio.* **12**, e0276621 (2021).
41. Yu, D., Xu, F., Valiente, J., Wang, S. & Zhan, J. An indigoidine biosynthetic gene cluster from *Streptomyces chromofuscus* ATCC 49982 contains an unusual IndB homologue. *J. Ind. Microbiol. Biotechnol.* **40**, 159-168 (2013).

42. Wang, L. et al. Protein scaffold optimizes arrangement of constituent enzymes in indigoidine synthetic pathway to improve the pigment production. *Appl. Microbiol. Biotechnol.* **104**, 10493-10502 (2020).
43. Xu, F., Gage, D. & Zhan, J. Efficient production of indigoidine in *Escherichia coli. J. Ind. Microbiol. Biotechnol.* **42**, 1149-1155 (2015).
44. Wehrs, M. et al. Correction to: Production efficiency of the bacterial non-ribosomal peptide indigoidine relies on the respiratory metabolic state in *S. cerevisiae. Microb. Cell Fact.* **18**, 218 (2019).
